# Quantitative Mapping of Sulfation, Iduronic Acid, and Secondary Structure in Glycosaminoglycans

**DOI:** 10.64898/2026.03.17.712318

**Authors:** Miguel Riopedre-Fernandez, Denys Biriukov, Hector Martinez-Seara

## Abstract

Glycosaminoglycans (GAGs) are extracellular matrix polysaccharides whose sequence variability and chemical modifications, particularly sulfation, generate substantial structural diversity. However, how sulfation patterns and monosaccharide composition encode secondary structure in GAGs is not systematically resolved, and quantitative metrics for classifying these structures are largely lacking. Here, we employ large-scale all-atom molecular dynamics simulations to investigate the molecular origin of secondary structure in sulfated GAGs. We systematically vary sulfation patterns and monosaccharide composition to isolate the factors that promote changes in three-dimensional structure. We show that GAG helical conformations arise from recurrent local shortening motifs caused primarily by stabilization of l-iduronic acid in the ^1^C_4_ puckering conformation, promoted by 2-O-sulfation or by densely sulfated regions. We also introduce a two-parameter structural metric that objectively classifies GAG secondary structures and distinguishes heparin helices from related conformations. Together, our results establish a quantitative link between monosaccharide identity, sulfation pattern, and three-dimensional organization of polysaccharide chains, providing a framework for future studies of sequence–structure relationships in GAGs.

## Introduction

Glycosaminoglycans (GAGs) are negatively charged polysaccharides that are widely distributed in the extracellular matrix (ECM) and connective tissues of many organisms.^1,2^ GAGs display remarkable structural diversity arising from variations in sequence and the presence of chemical modifications, among which sulfation plays a leading role. These chemical variations define the different types of GAGs and can vary substantially even within a single polysaccharide chain.^3,4^ Such structural heterogeneity often results in the organization of sulfated and nonsulfated monosaccharides into domains, commonly referred to as “sulfation patterns” or “sulfation code”.^5,6^ The sulfation pattern of GAGs plays a critical role in determining their interactions with proteins, ^7^ thereby influencing a wide range of cellular processes, including those associated with disease development.^2,3,5^ Despite extensive characterization, a systematic understanding of how specific sulfation patterns translate into defined secondary structures remains limited. Furthermore, the field lacks robust quantitative criteria to objectively identify secondary structure in flexible GAG chains.

Sulfation in GAGs profoundly influences the conformational behavior of their polysaccharide chains, thereby modulating their secondary structure. Secondary structures have been described for several GAGs, supported by numerous nuclear magnetic resonance (NMR), crystallographic, and molecular dynamics (MD) studies, some of which have resulted in deposited structures in the Protein Data Bank (PDB).^8–13^ However, these structures do not establish whether extended GAG chains adopt persistent secondary structures in solution, nor how such structures depend on sequence and sulfation pattern. For example, most available structures feature GAGs bound to proteins. In the few instances where unbound (free) GAGs have been structurally resolved, the fragments typically comprise no more than 12 monosaccharide units, and often fewer.^8,14^ Although such short oligosaccharides are informative about local conformations, they do not allow us to assess how these local features translate into the conformational behavior of extended polysaccharide chains.

Establishing a direct sequence–structure relationship in GAGs is experimentally challenging due to intrinsic heterogeneity in chain length and sulfation pattern. There have been efforts to control diversity by analyzing synthetic GAG oligomers with well-defined characteristics, but they are limited in size due to complex synthesis and purification.^15–18^ While NMR has been one of the key experimental techniques in saccharide research, ^19^ it provides only an ensemble-averaged picture of the accessible conformational space, making it challenging to effectively discriminate between multiple conformations that may coexist in solution.^20^ This situation is particularly relevant for large, flexible molecules like GAGs. Moreover, several reported “structures” are constructed from repeating subunits through computational refinement, potentially biasing the resulting models towards periodic and ensemble-averaged geometries.^8,21^ As a result, the available structures may not accurately represent a given polysaccharide chain’s true appearance.

The molecularly similar heparin (Hep) and heparan sulfate (HS) are distinguished from other GAGs by the presence of l-iduronic acid (IdoA) and their typically high negative charge density due to numerous sulfate and sulfamate groups (hereafter collectively referred to as “sulfate” groups). Hep and HS are reported to adopt helical-like conformations, often referred to as heparin helix (Hep–helix), depending on sulfation and sequence.^8,22,23^ The presence of the Hep–helix has been linked to the ability of Hep/HS to recognize and selectively bind specific proteins, such as fibroblast growth factors.^24^ Its occurrence is highly dependent on the puckering conformation of IdoA residues within the polysaccharide chain. ^24^ IdoA is a unique monosaccharide capable of adopting three distinct pucker conformations in solution: ^4^C_1_, ^2^S_0_, and ^1^C_4_,^22,25^ with sulfation affecting their relative populations. However, whether defined sulfation patterns together with monosaccharide sequence are sufficient to encode persistent secondary structures in extended GAG chains remains poorly understood. In this context, MD simulations provide a controlled framework to directly probe how defined sequences and sulfation patterns influence conformational ensembles over microsecond timescales.^17,26–29^

In this study, we employed large-scale all-atom MD simulations to characterize the secondary structures of relevant GAGs, primarily heparin/heparan sulfate (Hep/HS) and hyaluronan (HA). Our work explores the relationship between sequence, sulfation patterns, and three-dimensional structure. The studied systems include polysaccharide chains of up to 50 monosaccharides composed of relevant Hep/HS motifs^30^ or their analogs, allowing us to investigate both local and chain-wide effects of sulfation. As a control, we included HA, the only naturally occurring GAG that lacks sulfation and sequence variability,^31^ providing a baseline for assessing the extent of secondary structure in Hep/HS polysaccharides.

## Methods

### Simulated Systems

The summary of the simulated systems is provided in Tables 1 and S1 in the SI. Each system contained either 50- or 20-monosaccharide-long (50-mer and 20-mer, respectively) chains.

We constructed six large-scale systems (see Table 1 and Figure 1a for a graphical example) at three different saccharide concentrations: approximately 25 mM, 130 mM, and 260 mM, corresponding to 5, 25, and 50 polysaccharide chains, respectively. Three systems contained only hyaluronan (HA) chains, while the other three included a combination of two types of heparin (Hep) chains, Figure 1a–b. The first type of Hep chain (“NS”) contains three motifs characterized by a high concentration of sulfate groups and l-iduronic acid (IdoA) monosaccharides separated by repeating nonsulfated disaccharide sequences ([-*α*(1,4)-GlcA-*β*(1,4)-GlcNAc-]_n_), Figure 1c–d. The second type of Hep chain (“NANS”) features three motifs with a lower degree of sulfation. The NS and NANS sulfated domains have been suggested as fibroblast growth factor and antithrombin binding motifs, respectively.^4,30^

**Table 1:**
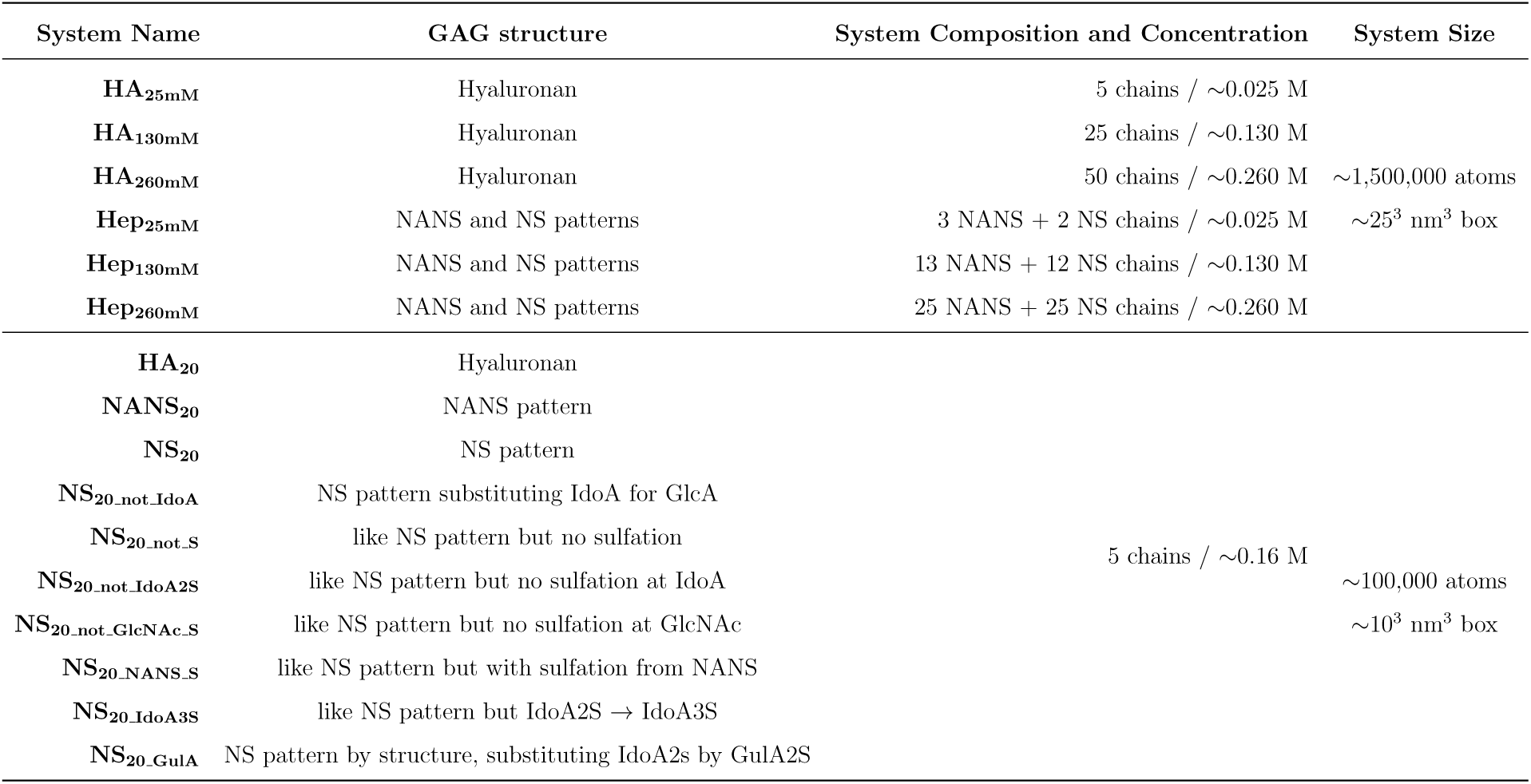
Summary of simulated systems, including GAG type, chain composition, saccharide concentration, and system size.

**Figure 1:**
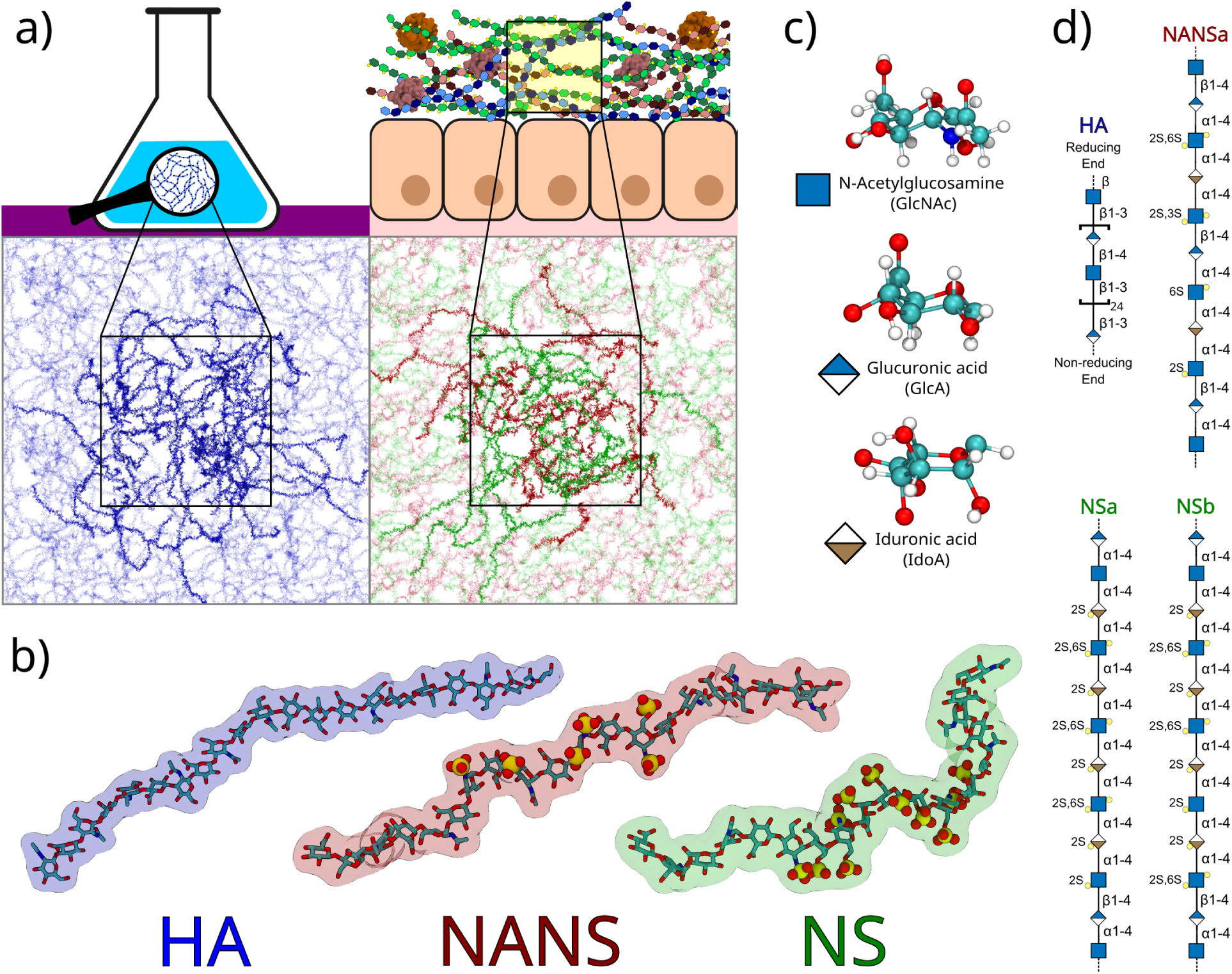
**a)** Schematic representation of biological contexts involving concentrated GAG solutions (synthetic hydrogel and glycocalyx) together with representative snapshots of HA_260mM_ and Hep_260mM_ simulation boxes. **b)** Representative conformations of central 20-mer segments of HA, NANS, and NS chains. **c)** Chemical structures of the monosaccharides used in this study: GlcNAc, GlcA, and IdoA. **d)** Symbol nomenclature for glycans (SNFG) illustrating HA and the sulfated domains of NANS (NANS_a_) and NS (NS_a_, NS_b_) chains.

Additionally, we prepared a set of ten smaller systems (Table 1, as well as Table S1 and Figure S4 in the SI) with custom sulfation patterns to identify the minimal components required for the formation of a GAG secondary structure. Each of these systems contained five 20-mer chains, corresponding to an approximate saccharide concentration of ∼0.16 M.

The structures of all polysaccharide chains are illustrated in Figures S1, S2, and S3 in the SI. The chains were built using the Glycan Reader & Modeler utility,^32^ available in the CHARMM-GUI online program.^33^ All systems were solvated with water, followed by the addition of ions to neutralize the net negative charge of GAGs and achieve a physiological NaCl concentration of ∼0.15 M.

As puckering changes occur on the microsecond timescale,^34^ the initial simulation setup of puckering in saccharides is critical. For this reason, we used ^1^C_4_ as the starting conformation for both IdoA and IdoA2S, as it has been reported to be the free energy minimum^25^ and the only distinguishable conformation after sulfation.^35^

### Simulation Models and Parameters

The all-atom prosECCo75 force field^36^ was used for all simulations in this study. The pros-ECCo75 force field is based on the CHARMM model^37^ and upgrades it by incorporating electronic polarization through charge-scaling in a mean-field approach. This force field includes all necessary parameters for simulating glycosaminoglycans,^38–41^ including the most recent parameters for sulfate groups in GAGs.^42^ In our prior research, we demonstrated that prosECCo75 outperforms the latest CHARMM-NBFIX model^37,43^ in modeling charged saccharides,^36,42,44,45^ while maintaining equivalent accuracy in other aspects, such as saccharide puckering and the distribution of glycosidic torsion angles.^36^ Parameters for sodium and chloride ions were adopted from the literature^46,47^ and incorporated into the prosECCo75 force field (labeled “Na s” and “Cl 2s”, respectively). Water was simulated using the TIP3P model.^48,49^

Simulations were conducted using the GROMACS simulation engine,^50^ versions 2021.3 and 2023.1. Buffered Verlet lists^51^ were used to maintain atomic neighbor lists. Long-range electrostatics was handled using the smooth particle mesh Ewald (PME) algorithm,^52^ with a direct cutoff automatically adjusted at 1.2 nm. The Lennard-Jones potential was used to model van der Waals interactions, with a cut-off of 1.2 nm and a gradual decrease of forces to zero starting at 1.0 nm. The Nosé–Hoover thermostat^53,54^ was applied separately to polysaccharides and solvent, maintaining a temperature of 310 K with a coupling time constant of 1 ps. Pressure was kept at 1 bar using the isotropic Parrinello–Rahman barostat,^55^ with a coupling time constant of 5 ps and compressibility of 4.5×10*^−^*^5^ bar*^−^*^1^. All covalent bonds involving hydrogen atoms were constrained using P-LINCS,^56,57^ while water geometry was constrained using SETTLE.^58^

Each system underwent energy minimization using the steepest descent algorithm, followed by a short equilibration of ∼100 ps. Production runs were 1 *µ*s long, with the first 100 ns excluded from analysis. The analysis was performed using GROMACS built-in tools, the MDAnalysis Python library,^59^ and in-house Python scripts.

### Simulation Analysis

#### Average inter-monosaccharide distance

The average inter-monosaccharide distance was calculated as a function of the number of monomers following a similar methodology as the one outlined by Kutálková et al..^60^ This metric reports the average chain extension as a function of monomer separation. We calculated the distance between the C1 atoms of monosaccharides separated by a specific number of monomers. The distance was then averaged across all possible combinations of the given separation and all simulation frames.

#### Local shortening analysis

For each residue, the C1–C1 distance to the residue three positions downstream was calculated. This distance was averaged across all equivalent chains and simulation frames. Other possible separations were also tested and are shown in the SI (Figure S6). The values obtained with this metric correspond to the lengths of every possible trisaccharide fragment of the chain, arranged by their position.

#### Contact analysis

Contact probabilities were defined as the fraction of frames in which a given monosaccharide formed at least one contact. Values were normalized by the number of equivalent monosaccharides across all chains. For an interaction to be considered, a heavy atom of the monosaccharide had to be within 5 Å from a heavy atom of a non-excluded monosaccharide residue. Residues from different chains and those directly connected by a glycosidic linkage were excluded.

#### Distribution of glycosidic dihedral angles

The distribution of Φ and Ψ dihedral angles, as defined by Φ =O_5_–C_1_–O_1_–C_X’_ and Ψ =C_1_– O_1_–C_X’_–C_X-1’_,^61^ was calculated for all possible glycosidic linkage types. X is the position 3 or 4 of the previous monosaccharide, depending on the type of glycosidic linkage (1–3 or 1–4).

#### Puckering analysis

Puckering Cremer-Pople angles ^62^ were calculated for every IdoA residue and classified as ^1^C_4_, ^2^S_0_ or ^4^C_1_ according to: ^1^C_4_: *θ >* 135*^◦^*; ^2^S_0_: 135*^◦^> θ >* 45*^◦^*; and ^4^C_1_: 45*^◦^ > θ* using an adapted version of the code provided by Balogh et al..^63^

#### Helical parameters calculation

Helical parameters were calculated following the method of Sugeta and Tatsuo Miyazawa.^64^ The HELANAL program^65^ was employed to calculate the helical parameters, as implemented in MDAnalysis.^66^ Although the MDAnalysis implementation is originally developed for protein helices, the formalism depends only on atomic coordinates and is therefore applicable to any polymer.

Glycosidic oxygens were used as the atom selection. For sulfated chains, helical parameters were computed separately for each sulfated domain. When fewer than nine atoms were available, the preceding glycosidic oxygen was included to satisfy algorithm requirements. For unsulfated polymers, central regions of comparable length were selected. Parameters were averaged across equivalent fragments and simulation frames.

To evaluate the robustness of the helical parameters against coordinate noise, synthetic helices were generated with and without added noise, see Figure S13 in the SI.

## Results and discussion

### Highly sulfated chains exhibit reduced extension

Highly charged polyelectrolytes are generally expected to exhibit increased persistence length due to electrostatic repulsion between charged monomers.^67,68^ However, our Hep/HS systems (hereafter Hep), composed of ∼1:1 mixtures of the relevant NANS and NS chains (see Figure 1), consistently display shorter end-to-end distances than HA, despite their substantially higher charge density, Figure 2. This trend is independent of GAG concentration.

**Figure 2:**
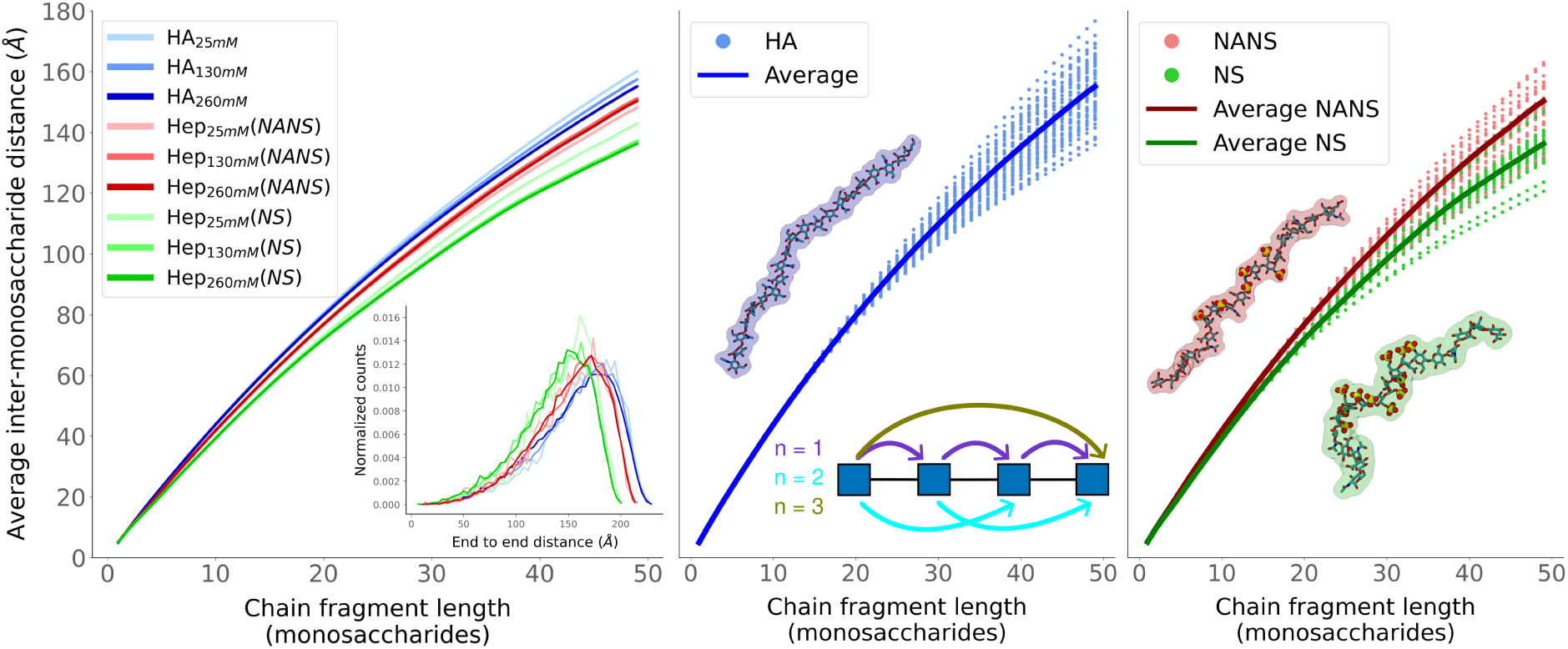
**Left:** Average C1–C1 inter-monosaccharide distance as a function of monomer separation for all 50-mer systems. The inset shows end-to-end distance distributions (equivalent to the distribution of the last points in the larger plot). Faster decay indicates increased chain bending and reduced persistence length. **Middle & Right:** Same metric shown separately for HA_260mM_ and Hep_260mM_, respectively. Solid lines represent the average over all chains; dashed lines correspond to individual chain averages. Representative chain conformations are shown for visual reference, along with a graphical representation of the metric.

The average inter-monosaccharide distance decays more rapidly with monomer separation in both NANS and NS chains compared to HA, indicating enhanced chain bending (Figure 2). The effect is strongest in NS chains, which possess the highest sulfation density. Because this metric directly relates to chain persistence length (Figure S5 in the SI), these results demonstrate that highly sulfated Hep chains are globally less extended than the less charged HA polymer.

This behavior contradicts simple electrostatics and suggests an underlying structural mechanism driving chain bending. A possible explanation is the formation of locally bent, kinked, or twisted structures. Given the sequence differences between HA and Hep, the bending likely arises from either iduronic acid (IdoA), specific sulfation patterns, or both. An alternative explanation could be that the reduced end-to-end distance arises from intrinsic differences in monosaccharide geometry or glycosidic bond lengths. However, analyses of glycosidic bond lengths show that HA single monosaccharide separations are comparable to, or smaller than, those in NS and NANS chains (see Figure S6 in the SI, panel 1). Therefore, the increased compactness cannot be attributed to monosaccharide-level geometric differences (see Figure S6 in the SI, panels 2–9).

### Local chain shortening is caused by iduronic acid and sulfation

To assess whether the global shortening arises from local structural features, we analyzed chain compactness using a fixed separation of three monosaccharides along the 50-mer chains (Figure 3, Top plot). Reduced distances in this metric indicate locally shortened regions. Notably, the sulfated regions of both NANS and NS chains, enriched in IdoA residues, display pronounced local shortening. These regions also exhibit an increased probability of intrachain contacts (Figure 3, Bottom plot). Both observations are consistent with the formation of compact structural motifs. The shortening in the sulfated regions is substantially greater in NS than in NANS, despite NS carrying five additional negative charges. This observation implies a stabilizing structural mechanism that locally outweighs Coulomb repulsion. In other words, instead of adopting extended conformations due to increased charge density, the highly sulfated NS segments preferentially fold into locally compact structures.

**Figure 3:**
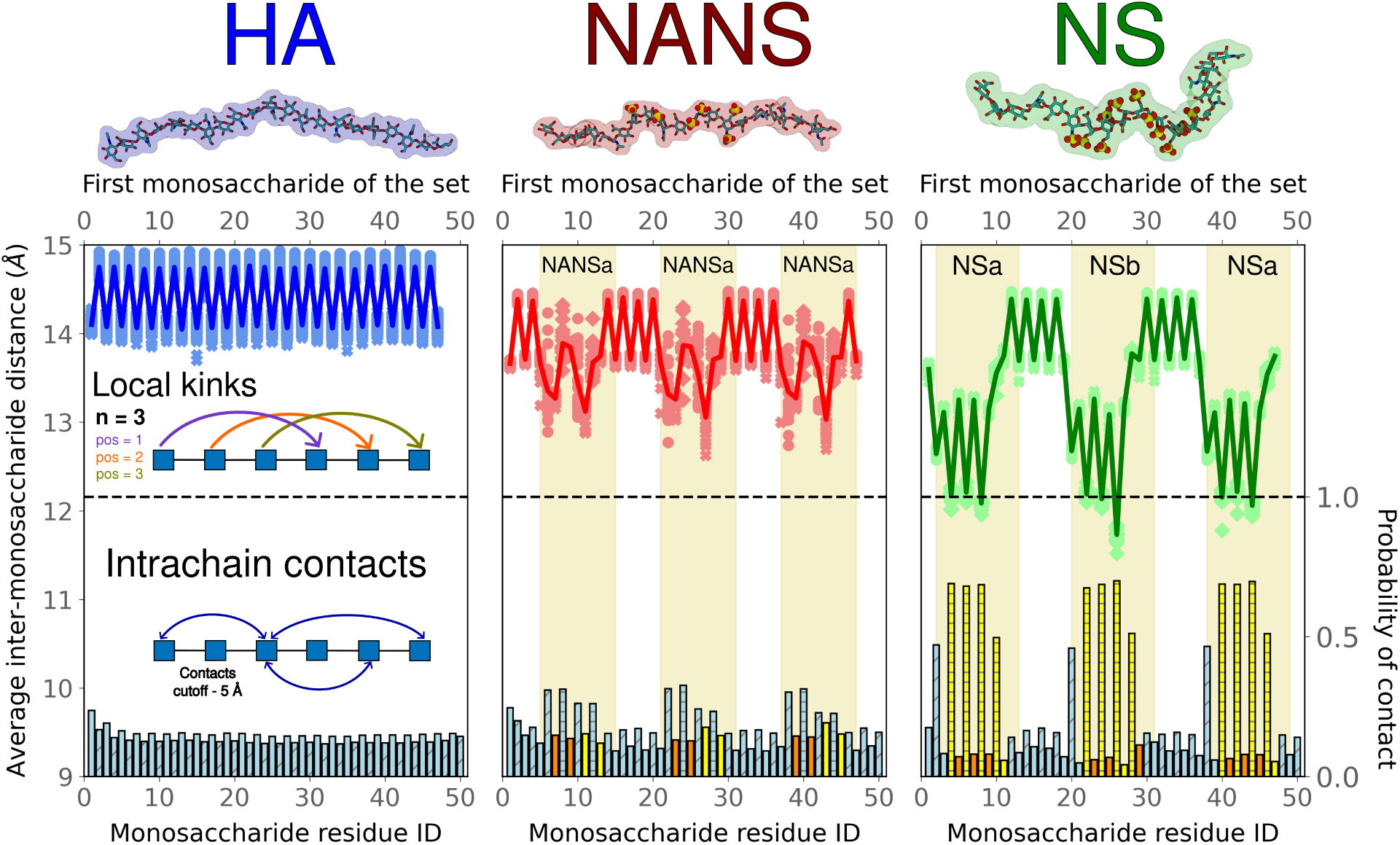
**Top plot:** C1–C1 distances between residues separated by three monosaccharides (*i* and *i* + 3) along HA_260mM_ and Hep_260mM_ chains. Reduced distances indicate local shortening. The solid lines indicate the average across all identical chains, while the dotted lines represent the average for each individual chain. The numbers on the *x*-axis indicate the number of the first monosaccharide of the trisaccharide. The yellow background boxes indicate the position and type of the sulfated regions (Figure 1d). The *y*-axis for this plot is on the left side. **Bottom plot:** Probability that each monosaccharide forms at least one intrachain contact (heavy-atom distance *<* 5^°^A). The monomers in the chains are numbered from one to fifty, with one being the reducing end and fifty the nonreducing end. Blue indicates a nonsulfated monosaccharide, yellow indicates a singly sulfated monomer, and orange indicates a monomer with two sulfate groups. Solid-filled bars represent GlcNAc; horizontal lines represent IdoA; diagonal lines represent GlcA. The *y*-axis for this plot is on the right side.

Further comparison between NS and NANS reveals reduced dispersion in the local distances of NS chains (points in Figure 3, Top plot), indicating a stable and reproducible secondary structure. In contrast, NANS chains exhibit broader variability, suggesting less defined conformational organization. While NANS displays some degree of local structuring in the sulfated regions, it appears insufficient to overcome electrostatic repulsion to the same extent as in NS. These results indicate that trisaccharides containing IdoA and sulfation constitute an optimal motif for inducing shortening, thereby altering secondary-structure formation.

HA behaves markedly differently. Its alternating disaccharide pattern produces regular distance profiles determined solely by trisaccharide identity (either GlcNAc–GlcA–GlcNAc or GlcA–GlcNAc–GlcA). Furthermore, intrachain contacts are largely absent, consistent with a stiffer and more homogeneous chain.

Overall, our data show that sulfation and IdoA drive local folding, which cumulatively leads to the overall chain shortening observed in Figure 2. We propose that the presence of these local deformations is what ultimately leads to the emergence of the Hep–helix. ^8^ The remaining question is therefore which molecular features control the folding.

To elucidate the molecular origin of this local folding, we measured lengths of the GlcNAc–IdoA–GlcNAc trisaccharides (C1*_i_* to C1*_i_*_+3_) as a function of different sulfation states, IdoA puckering conformations, and glycosidic dihedral angles. This motif was selected because it contains a central IdoA residue, which our data suggest plays a major role in shaping local chain geometry. The results shown in Figure 4 indicate that both puckering conformation and sulfation correlate with trisaccharide shortening. IdoA is naturally found in three puckering conformations: ^1^C_4_, ^4^C_1_, and ^2^S_0_. Both in NANS and NS, we find that the ^1^C_4_ conformer notably shortens the trisaccharide length. We attribute this to the effect of the ^1^C_4_ conformation on the overall geometry of the trisaccharide, which promotes zigzag and U-shaped conformations (see Figure S7 in the SI). The ^1^C_4_ conformation is typically the most prevalent state, with 2-O-sulfation further reinforcing this preference.^22,25,35^ In our simulations, unsulfated IdoA, present in the NANS patterns, readily transitions among all three available conformers. Instead, IdoA2S, which is characteristic of NS chains, remains predominantly in the ^1^C_4_ conformer throughout the simulations, consistent with experimental data.^25,35^ However, puckering transitions occur on the microsecond timescale;^34^ therefore, our systems may not have sampled less probable yet accessible states. Importantly, our conclusions do not rely on quantitative equilibrium populations, but on the structural consequences of stabilizing a specific puckering state.

**Figure 4:**
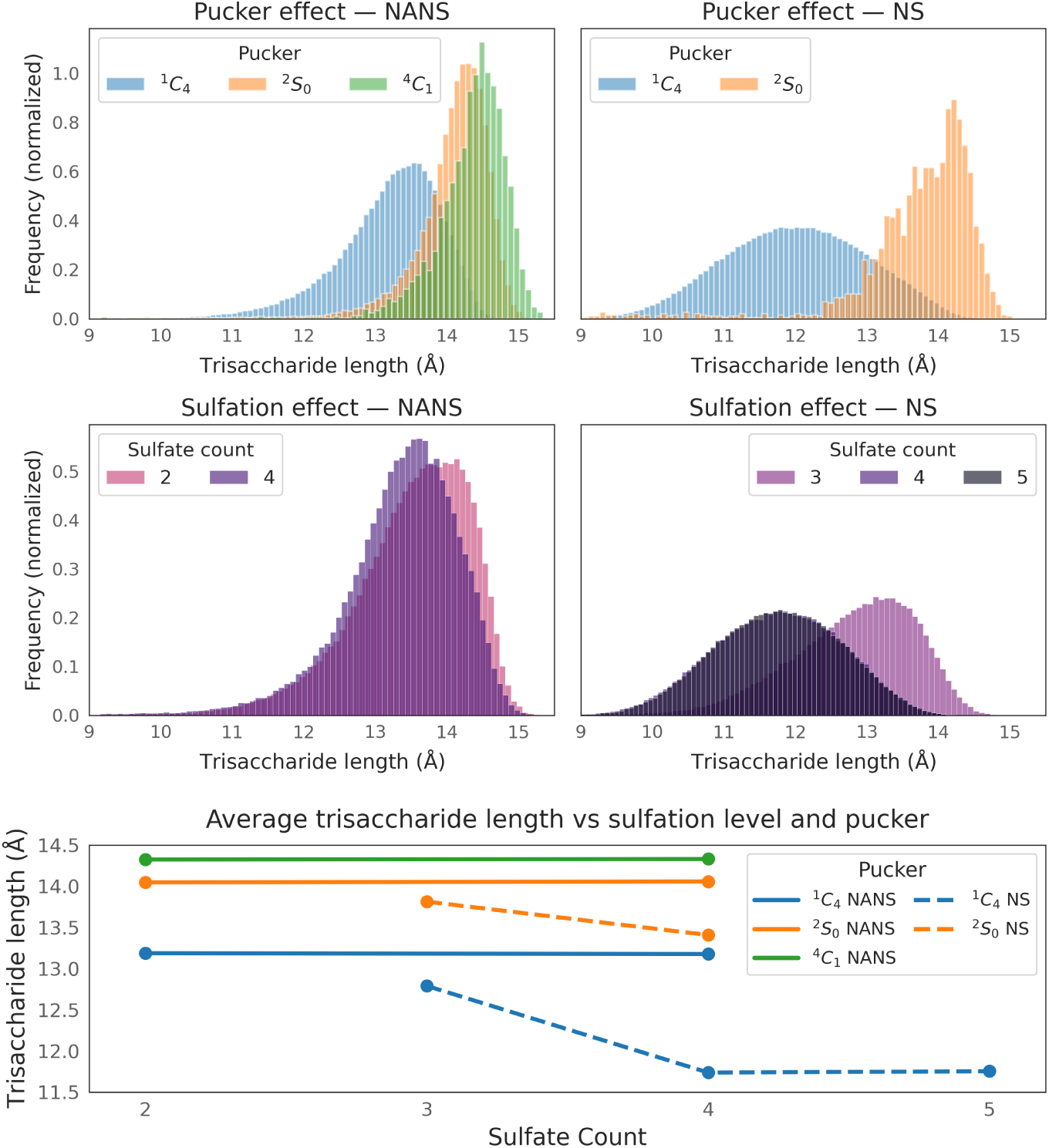
Distributions of the length of GlcNAc–IdoA–GlcNAc trisaccharides in the Hep systems, as a function of **Top:** IdoA pucker conformation (^1^C_4_, ^2^S_0_, and ^4^C_1_); and **Middle:** Number of sulfate groups in the trisaccharide. The distributions are shown separately for NANS (left) and NS chains (right). NANS contains IdoA, whereas NS contains IdoA2S. Histograms are normalized to unity; therefore, the curves represent probability densities rather than the absolute number of samples. **Bottom:** Average trisaccharide length as a function of both sulfate count and IdoA pucker conformation. NANS is shown as solid lines, while NS is represented by dashed lines. Standard errors of the mean are smaller than the markers. Notably, trisaccharides containing five sulfate groups never adopt puckering states other than ^1^C_4_.

One might assume that IdoA2S sulfation alone is sufficient to locally shorten a Hep chain by increasing the ^1^C_4_ population. However, trisaccharides containing isolated IdoA2S are, on average, more extended than those located in NS densely sulfated regions (Figure 4, Middle-Right and Bottom). Increasing sulfation affects only NS trisaccharides, indicating that local charge density plays a key role in reducing trisaccharide length. Furthermore, increasing the number of sulfate groups from four to five does not lead to additional shortening, as all three monosaccharides are already sulfated in both cases. Together, these findings indicate that the combination of sulfation pattern in the trisaccharide and IdoA puckering modulates chain shortening.

We also examined whether backbone glycosidic geometry could account for the observed differences in chain compaction. Unlike proteins, where dihedral angles largely determine local environments and ultimately secondary structure, the role of glycosidic bonds in these GAGs appears less significant. The dihedral distributions of glycosidic linkages are remarkably similar between HA, NANS, and NS chains (Figures S8, S9, S10 in the SI), with only a slightly narrower distribution in highly sulfated chains.

Overall, our results demonstrate that 2-O-sulfation of IdoA plays a dual role in Hep structuring: it biases the ring toward the shortening-prone ^1^C_4_ conformation and contributes to the formation of densely sulfated domains that amplify local compaction. The accumulation of these shortened motifs generates recurrent regions with distinct local folding along the chain (Figure 3).

Importantly, the relationship between these local folding motifs and the emergence of helicity is not straightforward. In the following sections, we dissect the structural descriptors underlying local folding and examine their connection to the Hep–helix and other secondary structures.

### Custom systems: building blocks of a helical Hep

It is interesting that the ^1^C_4_ conformation has been shown to promote helical structures in algae polysaccharides,^13^ although in the absence of IdoA. This could be interpreted as convergent evolution, in which different organisms address a potential structural requirement for helical saccharides through somewhat similar yet distinct mechanisms. To identify the minimal structural requirements for the formation of local folding motifs leading to the Hep–helix or other secondary structures, we prepared an additional set of ten systems containing 20 monosaccharide-long polysaccharide chains (Tables 1 and S1 in SI) in which the presence, position, and type of IdoA residues and sulfate groups were systematically varied. These controlled modifications allowed us to test which molecular features are necessary and/or sufficient to induce local folding. A comprehensive summary of the custom systems is provided in the SI (Figures S2, S3, S4, and Table S1).

For HA, NS, and NANS, the 20-mer systems in Figure 5 show very similar behaviors to their longer counterparts in Figure 3. Interestingly, NS_20_ exhibits one of the shortest end-to-end distances among the custom sequences (Figure S11 in the SI), consistent with the accumulation of regularly spaced bending motifs (Figure 5). As this corresponds to a naturally occurring sulfation pattern, it suggests that the distinctive behavior of this sequence may be functionally relevant. In fact, consistent with this interpretation, we observe clear patterns in the behavior of the custom fragments depending on the presence and distribution of IdoA residues and sulfation motifs (Figure 5).

**Figure 5:**
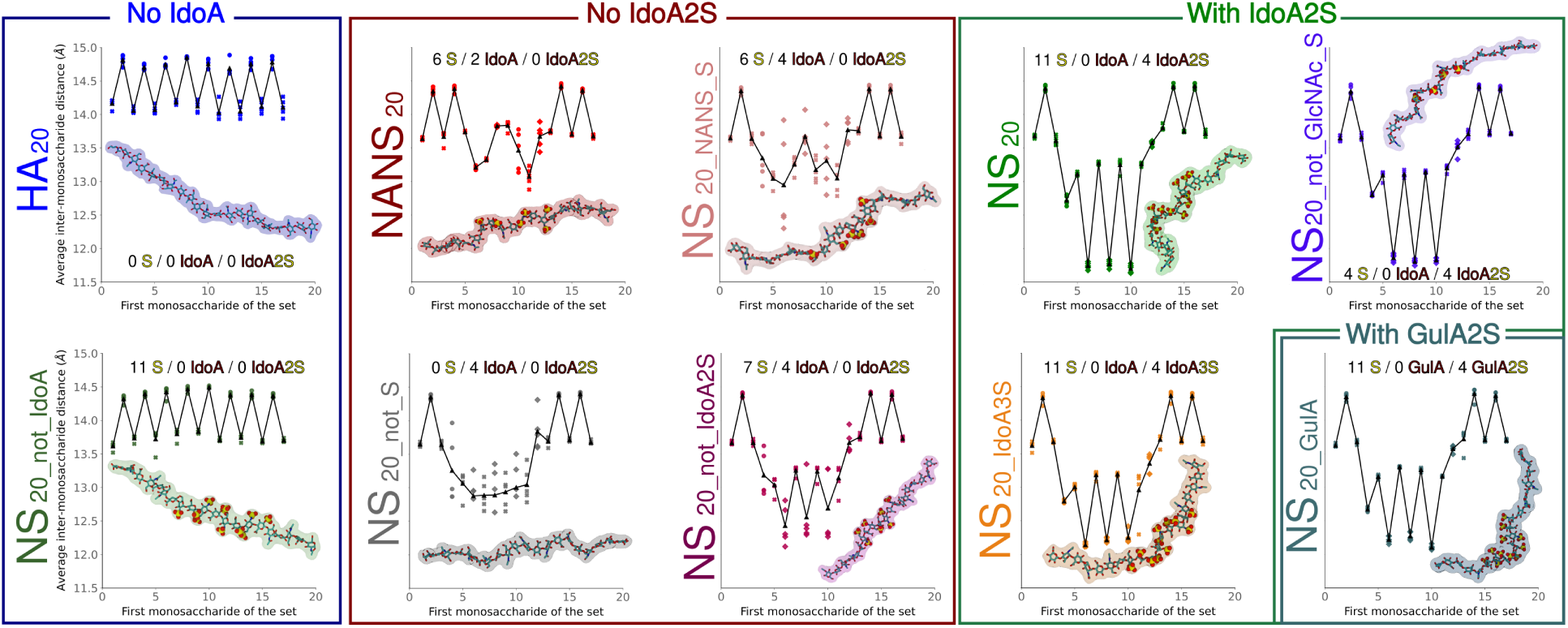
Average C1–C1 distances for residues separated by three monosaccharides (solid lines) are shown for each system. Individual chain averages are shown as points. Systems are grouped based on their IdoA, IdoA2S, or GulA2S content. The number of sulfate groups, IdoA residues, and IdoA2S residues per chain is indicated.

NS chains in which IdoA is replaced by GlcA show a regular local shortening pattern, comparable to HA (Figure 5, “No IdoA”), indicating the absence of localized bending and reflecting only intrinsic differences between repeating trisaccharides. This confirms IdoA as a necessary component for generating deviations from regular patterns and extended topologies.

Sequences containing IdoA show shortening in the areas adjacent to IdoA residues, although the effect is generally weaker than in systems with IdoA2S (Figure 5, “No IdoA2S” vs. “With IdoA2S”). However, sequences with IdoA exhibit a higher degree of structural variability, as evidenced by the broader distribution of individual average chain lengths (dots in Figure 5, each representing a different chain). This variability arises from the flexibility of the IdoA ring and its ability to interconvert between puckering conformations in our simulations.

Compared to all other studied sequences with only IdoA, NS_20 not IdoA2S_, which preserves the NS sequence and most of its sulfations except those in IdoA2S, exhibits a shortening pattern similar to, but weaker than, that of NS_20_. This indicates that sulfation of neighboring GlcNAc residues partially stabilizes local bending and compacting even when the IdoA conformation is not locked, further evidencing the structuring role of dense sulfation environments. Nevertheless, the overall structural heterogeneity remains higher than in fully sulfated NS systems.

The introduction of IdoA2S systematically produces a characteristic jagged profile that closely resembles the NS pattern (Figure 5, “With IdoA2S”). The narrow and periodic distribution indicates a stable pattern of localized bends along the chain.

Interestingly, removing the sulfate groups from GlcNAc (NS_20 not GlcNAc S_) produces a similar shortening pattern to NS_20_ if not slightly more pronounced. This observation is consistent with this system exhibiting the shortest end-to-end distance (Figure S11 in the SI). These results suggest that IdoA2S alone is sufficient to induce shortening by stabilizing the ^1^C_4_ conformation. In this context, removing neighboring sulfations likely reduces electrostatic repulsion within the cluster of negatively charged sulfate and carboxylate groups, further enhancing chain shortening without altering the puckering equilibrium.

To explore the importance of 2-O-sulfation in IdoA, we prepared a system containing IdoA with 3-O-sulfation (IdoA3S). Although IdoA3S is not present in mammals, it has recently been identified in halophilic archaea.^69^ In our simulations, IdoA3S remains mostly in the ^1^C_4_ conformer. This system exhibits a similar behavior to IdoA2S with respect to localized shortening, suggesting that stabilization of the ^1^C_4_ conformation, rather than the precise sulfate position, is the dominant factor controlling local bending. This finding is consistent with reports that algal polysaccharides^13^ with monosaccharides fixed to ^1^C_4_ conformation via intermolecular bridging display helical folding.

To verify this logic, we generated a Hep-like sequence in which IdoA2S was replaced by 2-O-sulfated guluronic acid (GulA2S). This non-natural monosaccharide has its two bulky substituents (carboxylate and sulfate) in equatorial positions in the ^1^C_4_ conformation, which further stabilizes this state. In our simulations, no conformational transitions were observed, effectively fixing the ring in the ^1^C_4_ state. This system exhibits pronounced localized shortening and global chain bending, comparable to those observed in the other IdoA2S-containing polysaccharide chains tested. However, despite visually displaying a helical structure, the overall structure is distinct from NS_20_, which is more similar to the canonical Hep–helix.^8^ This demonstrates that stabilization of the ^1^C_4_ conformation generates helicity-like behavior, but this alone is not sufficient to reproduce a specific secondary structure.

Together, these results demonstrate that IdoA is necessary in Hep to induce localized bending motifs along the chain but is not sufficient to produce a stable and compact bent structure. The presence of IdoA2S is sufficient to generate stable and reproducible local folds, while dense sulfation environments around IdoA can partially stabilize shortening patterns. We finally generalize that monosaccharides and sulfation patterns that stabilize the ^1^C_4_ conformation may promote helical structures, although not necessarily identical to canonical Hep–helices.^8^ This observation underscores the need to develop an objective metric capable of distinguishing secondary structures that emerge from the observed local patterns.

### Objective determination of GAG secondary structures

The fact that similar shortening patterns can produce different bent structures highlights the need for an objective metric to quantify secondary structure in our systems and in GAGs in general. Defining the secondary structure, including the experimentally described helicity,^8^ of a highly variable three-dimensional polymer is inherently challenging. Existing strategies, *e.g.*, those for *α*-helices in proteins, are often not applicable to GAGs. This is evidenced by the limited usefulness of glycosidic dihedral angle distributions (Figures S8, S9, S10 in the SI), which contrasts with the clear interpretability of Ramachandran plots for proteins.

Focusing on helicity, the formalism introduced by Sugeta and Tatsuo Miyazawa ^64^ enables the calculation of helical parameters for an arbitrary polymer chain. The remaining challenge is to determine which combination of these parameters effectively discriminates between helical and non-helical structures. Previous studies have primarily used helical pitch as a sole descriptor of helicity,^13^ but this metric alone does not capture the full complexity of GAG conformations.

We found that two parameters, the local twist angle (*i.e.*, the angular displacement between consecutive residues) and the number of residues per turn, clearly distinguish our different GAG structures (Figure 6). Chains enriched in IdoA2S, IdoA3S, and densely sulfated IdoA environments cluster in the upper-left region of the plot. Notably, all chains displaying localized shortening and recurrent bending motifs fall within this well-defined region, with the exception of NS_20 GulA_. Interestingly, this is the region where the Hep–helix from PDB:1HPN,^8^ with all IdoA2S residues in ^1^C_4_, appears after a short MD simulation. This allows us to define this region as characteristic of the Hep–helix secondary structure.

**Figure 6:**
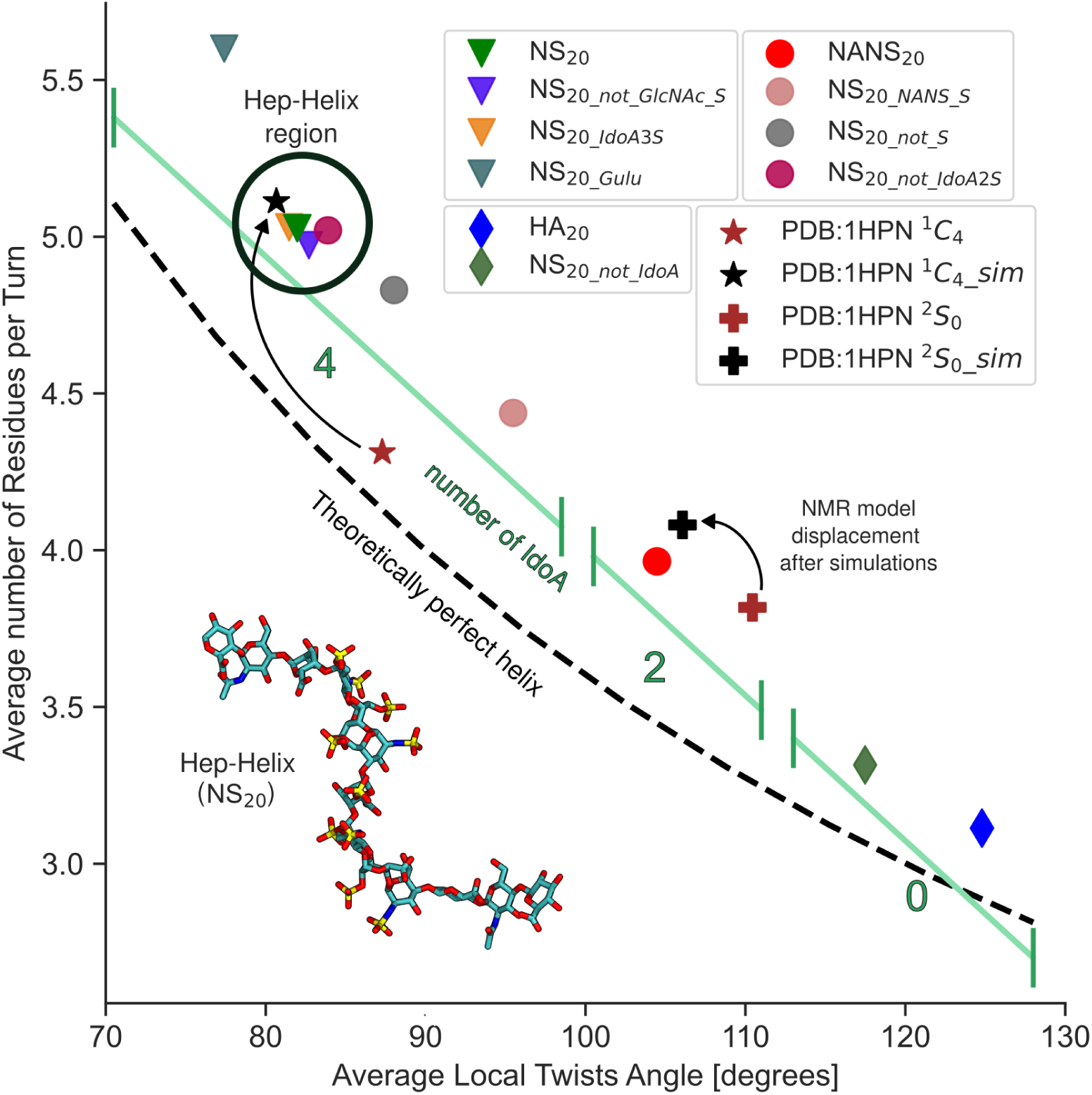
Average local twist angle and average number of residues per turn calculated using the Sugeta and Tatsuo Miyazawa formalism.^64^ Only the data from 20-mer systems are shown; 50-mer systems can be seen in Figure S12 in the SI. Each point represents a sulfated domain (NS, NANS) or equivalent region in HA. The dashed black line corresponds to the geometric ideal helix (twist × residues per turn = 360°). Chains enriched in IdoA and IdoA2S cluster within a well-defined Hep–helix region in the top-left. The GulA-containing system lies outside this cluster despite enforced ^1^C_4_ puckering. The values from the static NMR-derived PDB:1HPN structure and MD simulations starting from this model, both restrained and unrestrained, are included for comparison.

At the opposite end of the parameter space, HA and NS systems lacking IdoA occupy the bottom-right region. Furthermore, the IdoA content correlates with the position on the metric (Figure 6, green line). Therefore, this two-parameter representation appears to provide a quantitative, objective method for classifying GAG structures in solution, including their helicity and structural relationships. In this representation, the GulA-containing chain occupies the extreme upper-left region of the plot, reflecting its strongly enforced ^1^C_4_ conformation. However, it lies outside the canonical Hep–helix cluster, confirming that stabilization of ^1^C_4_ alone does not guarantee a Hep–helix topology, and thereby validating the discriminatory power of the metric.

To further evaluate the method, we analyzed the two NMR-derived structures of the Hep dodecamer (PDB:1HPN^8^). The PDB entry contains two structures: one with all IdoA2S in the ^1^C_4_ conformation introduced above, and the other in the ^2^S_0_ conformation. We applied our secondary structure analysis to both restrained and unrestrained MD simulations starting from these two models. In the restrained simulations, puckering transitions were prevented (Figure 6). Contrary to the ^1^C_4_ model, which moved to the Hep–helix region, the ^2^S_0_ model starts in a different region and shifts toward conformations characteristic of NANS-like chains. This result further illustrates the conformational heterogeneity of flexible GAGs.

The metric remains consistent across both 20-and 50-mer systems and across different concentrations (Figure S12 in the SI), demonstrating its stability with respect to system size and sampling conditions. Taken together, this framework provides a quantitative link between sequence, IdoA content, sulfation pattern, and three-dimensional topology. It offers a practical tool for distinguishing related but structurally distinct helical arrangements and may potentially predict whether a given GAG sequence is capable of forming specific secondary structures.

## Conclusions

This work elucidates the mechanisms underlying secondary structure formation in biologically relevant glycosaminoglycans (GAGs), namely heparin/heparan sulfate (Hep) and hyaluronan. To this end, we performed a set of large-scale molecular dynamics simulations to identify the key modifiers of GAG structure. We find that IdoA-rich and highly sulfated chains are more compact than their less charged counterparts, despite their higher charge density. This demonstrates that specific conformational preferences can overcome electrostatic repulsion and govern the local folding of GAGs.

We identify the locking of IdoA into its ^1^C_4_ conformation as the main molecular mechanism underlying local chain shortening. 2-O-sulfation (IdoA2S) biases the ring toward this shortening-prone geometry,^25,35^ thereby promoting regular folds along the chain. In addition, densely sulfated environments can further enhance local compaction. We connect these local motifs to the emergence of secondary structures, which remain poorly defined in the GAG field.

To address this gap, we introduce an objective two-parameter metric based on the local twist angle and the number of residues per turn that distinguishes among different GAG helices and other conformations. This representation links GAG sequence, ring puckering, and sulfation patterns to their position in the metric and the resulting secondary structure. Within this representation, GAGs enriched in IdoA2S and sulfate groups cluster around a specific region associated with the proposed heparin helix. We anticipate that this metric will provide a quantitative reference for future studies aiming to characterize and compare GAG secondary structures. Further work on the role of secondary structure in GAG bioactivity will be essential to understand how biological systems tune GAG function through local structural changes and to guide the design of bioactive GAGs.

## CRediT authorship contribution statement

**Miguel Riopedre-Fernandez:** Conceptualization, Methodology, Investigation, Formal analysis, Visualization, Writing – Original Draft.

**Denys Biriukov:** Conceptualization, Methodology, Resources, Supervision, Project administration, Writing - Review and Editing.

**Hector Martinez-Seara:** Conceptualization, Methodology, Resources, Supervision, Project administration, Writing - Review and Editing.

## Supporting information

The Supporting Information includes the complete sequences of the simulated GAG chains, structures of the 20-mer systems, and additional molecular dynamics analyses.

## Data availability

All simulation data and the helical parameters code have been uploaded to the public scientific repository Zenodo (DOI: 10.5281/zenodo.19054422).

## Acknowledgements

M.R.-F. acknowledges the support from Charles University in Prague and the International Max Planck Research School in Dresden.

D.B. acknowledges VSB – Technical University of Ostrava, IT4Innovations National Supercomputing Center, Czech Republic, for awarding this project access to the LUMI supercomputer, owned by the EuroHPC Joint Undertaking, hosted by CSC (Finland) and the LUMI consortium through the Ministry of Education, Youth and Sports of the Czech Republic through the e-INFRA CZ (grant ID: 90254), project OPEN-35-3.

The authors acknowledge Grammarly and ChatGPT for improving the readability and language of the manuscript.

