## Supplementary Information for "Quantitative Mapping of Sulfation, Iduronic Acid, and Secondary Structure in Glycosaminoglycans"

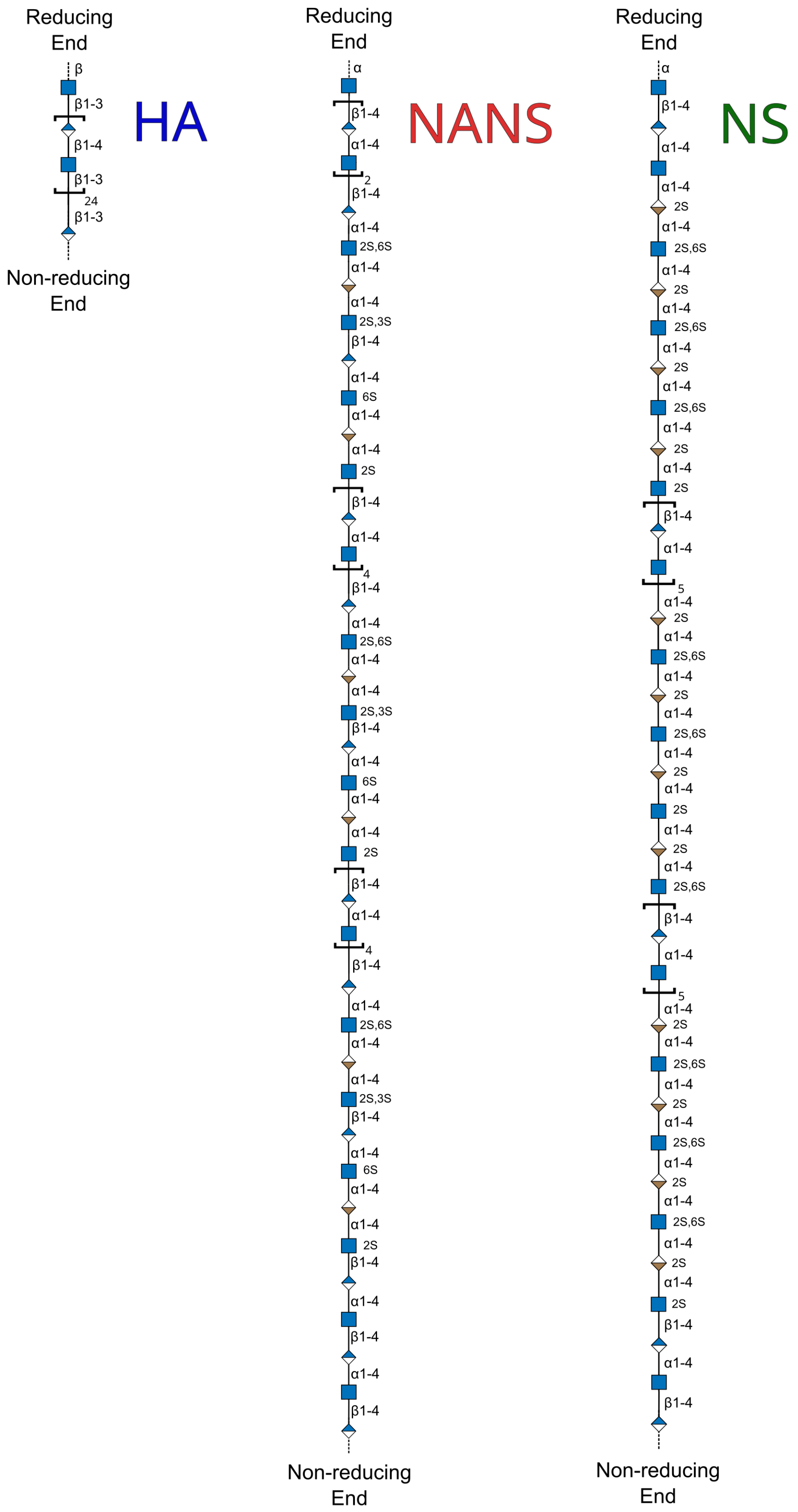


Figure S1: Symbol Nomenclature for Glycans (SNFG) representation of the polysaccharide chains used in the 50-mer simulations. The structures of hyaluronan (HA) and the two heparin motifs (NANS and NS) are shown from the reducing to the non-reducing end.

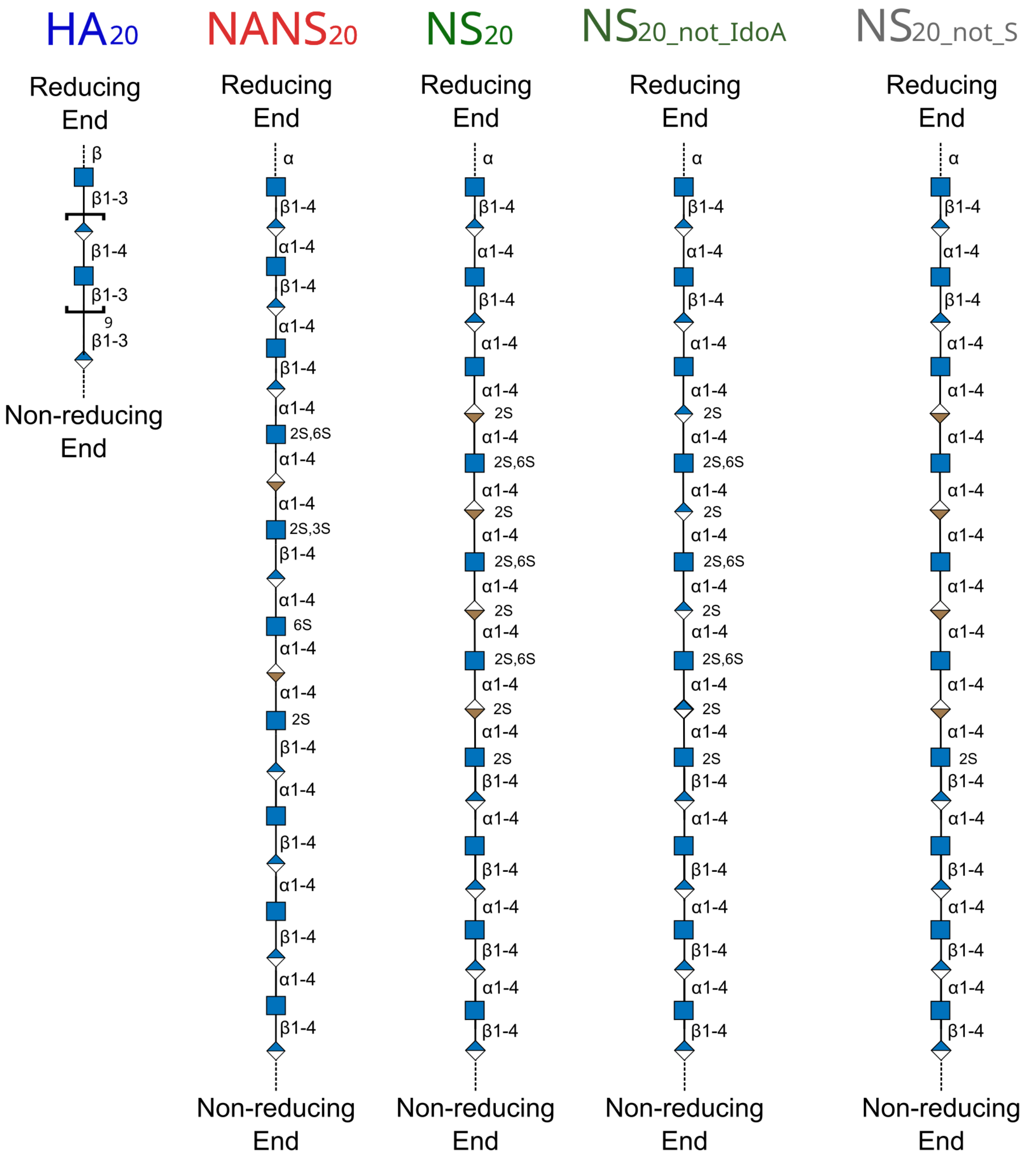


Figure S2: SNFG representation of five 20-mer systems: $HA_{20}$, $NANS_{20}$, $NS_{20}$, $NS_{20\_not\_IdoA}$, and $NS_{20\_not\_S}$.

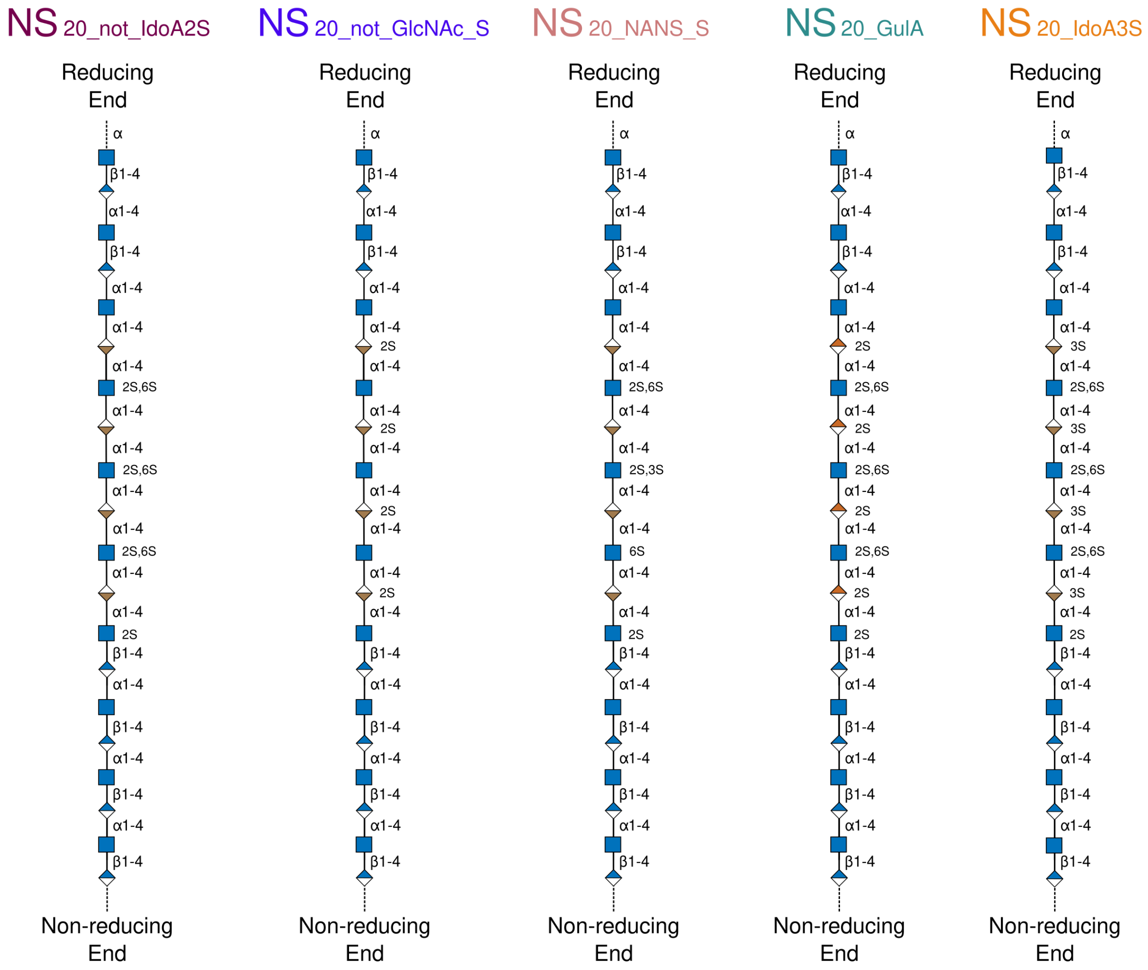


Figure S3: SNFG representation of the other 20-mer systems not shown in Figure S2: $NS_{20\_not\_IdoA2S}$, $NS_{20\_not\_GlcNAc\_S}$, $NS_{20\_NANS\_S}$, $NS_{20\_GulA}$ and $NS_{20\_IdoA3S}$.

Table S1: Summary of the custom 20-mer systems used to dissect the structural determinants of helicity. For each system, the table lists the design rationale, the total number of sulfate groups, iduronic acid residues (IdoA), 2-O-sulfated iduronic acid residues (IdoA2S), and other relevant residues per chain, along with the presence or absence of helical structure observed in the simulations.

| **System Name** | **Explanation** | **Number of sulfates / IdoA / IdoA2S / other per chain** | **hep–Helix observed** |
|---|---|---|---|
| $\mathbf{HA_{20}}$ | HA | 0 / 0 / 0 | No |
| $\mathbf{NANS_{20}}$ | NANS pattern | 6 / 2 / 0 | No |
| $\mathbf{NS_{20}}$ | NS pattern | 11 / 0 / 4 | Yes |
| $\mathbf{NS_{20\_not\_IdoA}}$ | NS pattern substituting IdoA for GlcA | 11 / 0 / 0 | No |
| $\mathbf{NS_{20\_not\_S}}$ | NS pattern removing all sulfations | 0 / 4 / 0 | No |
| $\mathbf{NS_{20\_not\_IdoA2S}}$ | NS pattern removing the sulfations in IdoA | 7 / 4 / 0 | Yes |
| $\mathbf{NS_{20\_not\_GlcNAc\_S}}$ | NS pattern removing the sulfations in GlcNAc | 4 / 0 / 4 | Yes |
| $\mathbf{NS_{20\_NANS\_S}}$ | NS pattern by structure, with sulfation from NANS | 6 / 4 / 0 | No |
| $\mathbf{NS_{20\_IdoA3S}}$ | NS pattern substituting the IdoA2S for IdoA3S | 11 / 0 / 0 / 4 IdoA3S | Yes |
| $\mathbf{NS_{20\_GulA}}$ | NS pattern by structure, substituting IdoA2s by GulA2S | 11 / 0 / 0 / 4 GulA2S | Other Helix |

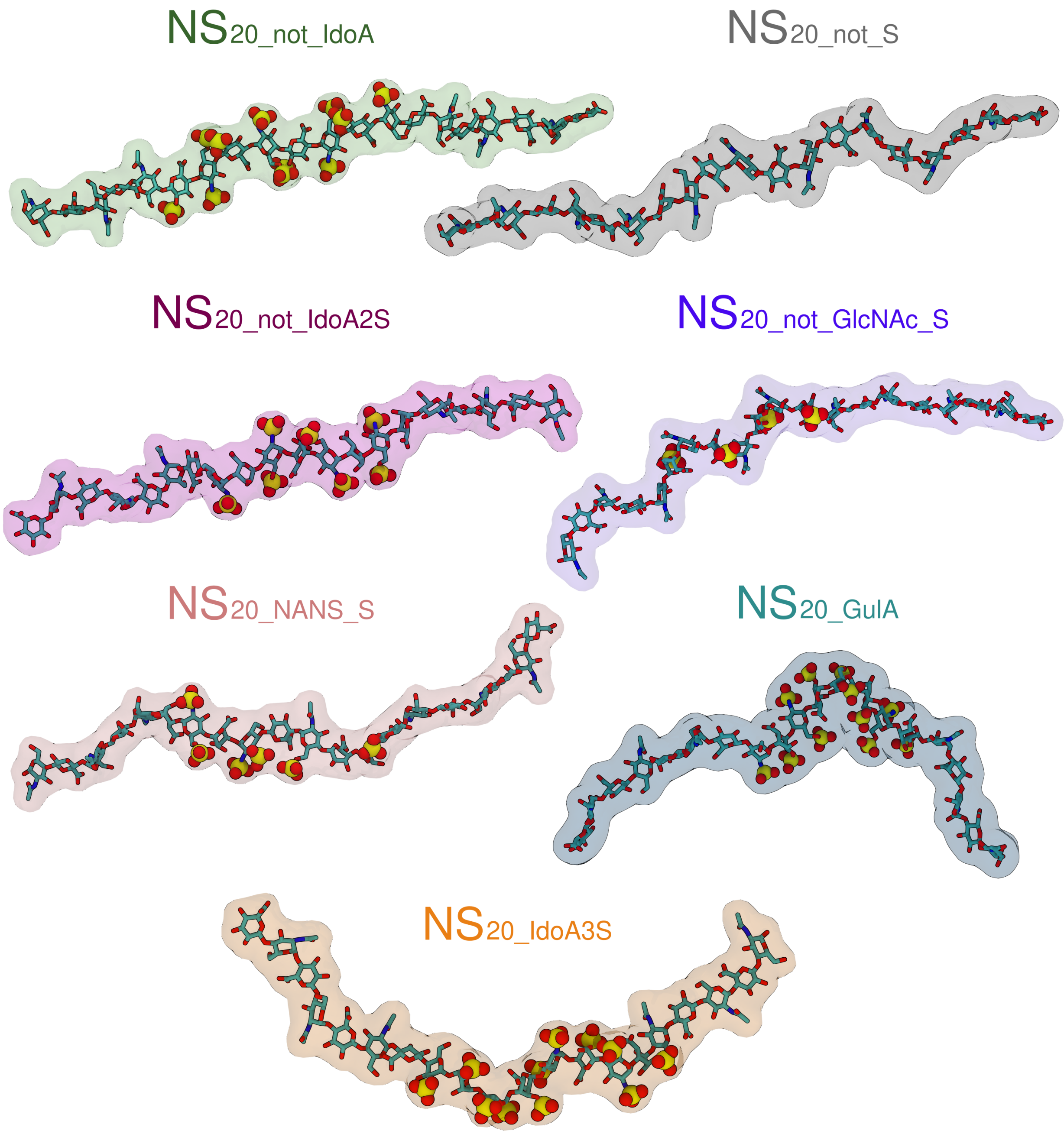


Figure S4: Representative conformations of the 20-mer systems extracted from the simulations.

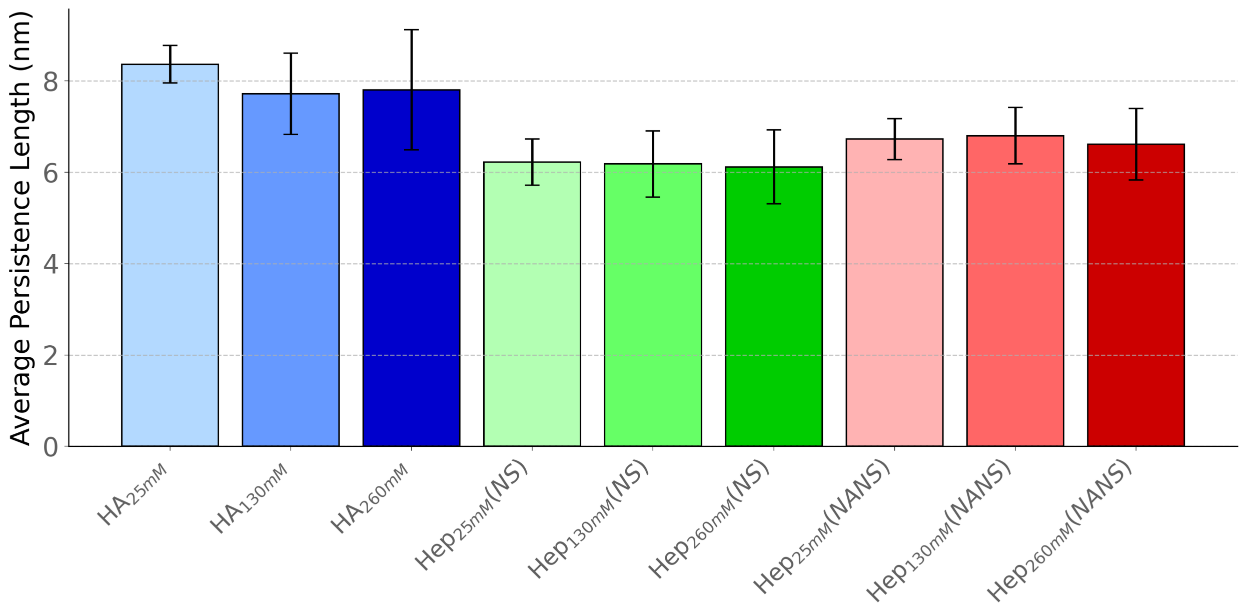


Figure S5: Average persistence length of the polysaccharide chains in the 50-monosaccharide systems. Persistence lengths were calculated from the decay of the tangent–tangent correlation function along the polymer backbone. Tangent vectors were defined between consecutive C1 atoms, and the correlation $\langle \cos\theta_i \rangle$ between normalized tangent vectors separated by $i$ bonds was averaged over all simulation frames and chains. The resulting decay was fitted to the worm-like chain relation $\langle \cos\theta_i \rangle = e^{-i/L_p}$, where $L_p$ is the persistence length expressed in bond units. The persistence length in physical units was obtained by multiplying $L_p$ by the average distance between consecutive C1 atoms. Bars represent mean persistence lengths for each system, and error bars correspond to the standard deviation across individual chains.

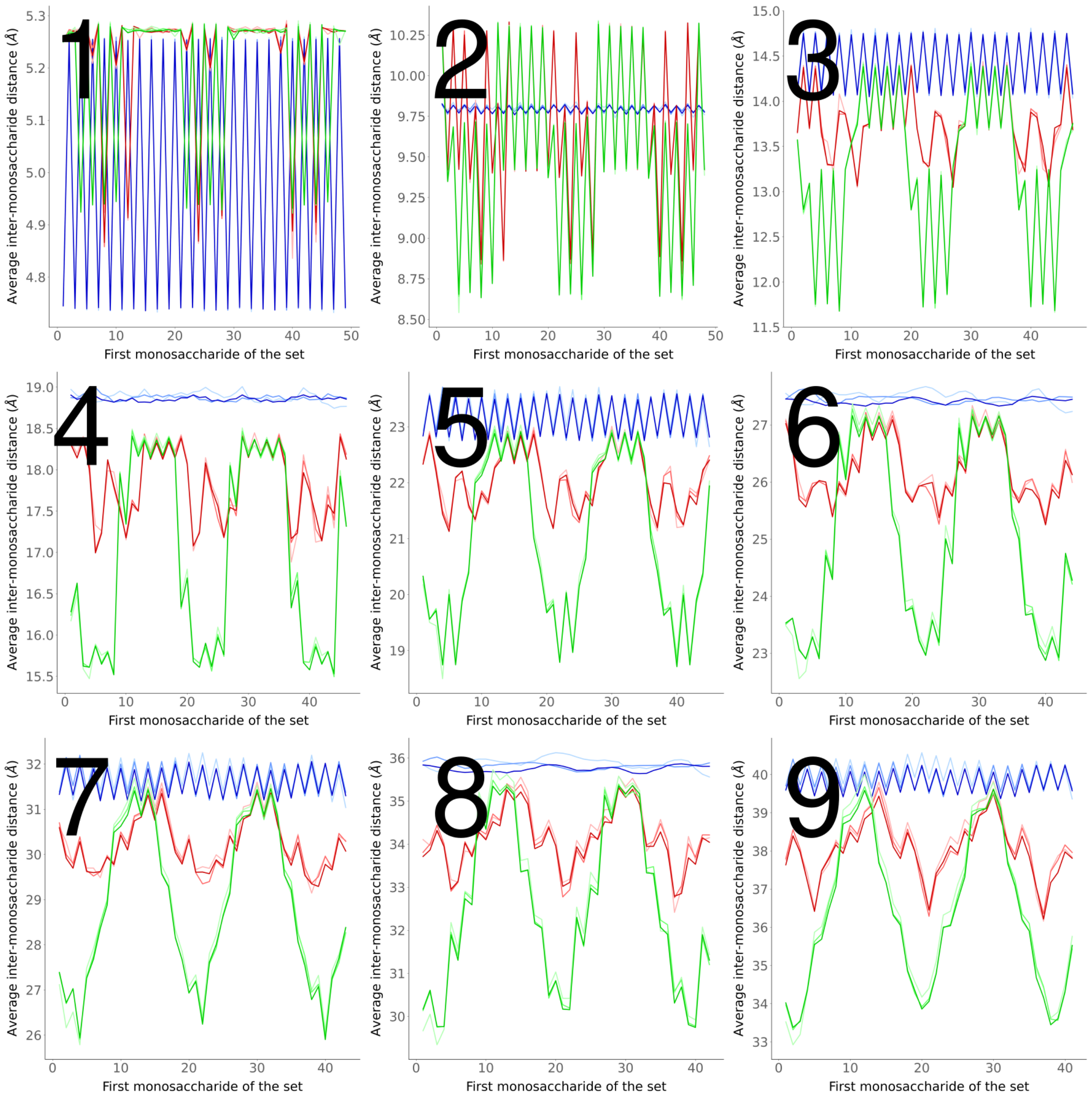


Figure S6: C1–C1 distances between residues separated by one to nine monosaccharides along the backbone in all 50-mer systems. Blue lines represent HA chains, red lines are NANS chains, and green lines are NS chains. Color intensity reflects polysaccharide concentration, with concentrations ranging from 0.25 mM to 260 mM represented by increasing color darkness. Shorter distances indicate localized chain shortening. The numbers on the $x$-axis indicate the index of the first monosaccharide in the fragment used for the distance calculation.

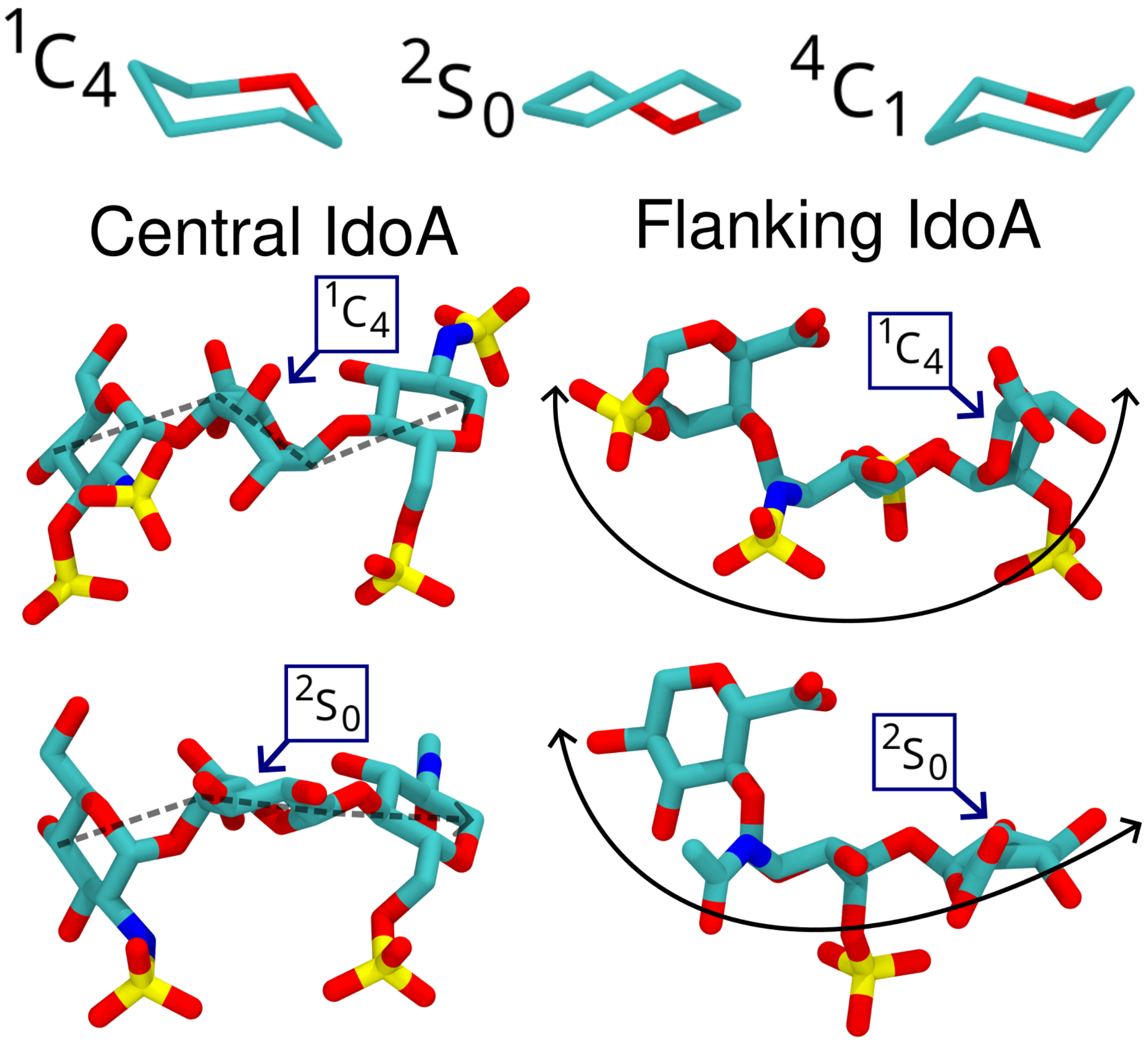


Figure S7: Graphical representation of IdoA mono- and trisaccharides in different puckering conformations. **Top:** IdoA monosaccharides in the $^{1}C_{4}$, $^{2}S_{0}$, and $^{4}C_{1}$ conformations. **Middle & Bottom:** Two types of trisaccharides are shown: those with a central IdoA residue (GlcNAc–IdoA–GlcNAc) and those with flanking IdoA residues (IdoA–GlcNAc–IdoA or GlcA–GlcNAc–IdoA). Dashed arrows highlight the approximate plane of the pyranose ring, while solid arrows illustrate the overall curvature of the trisaccharide.

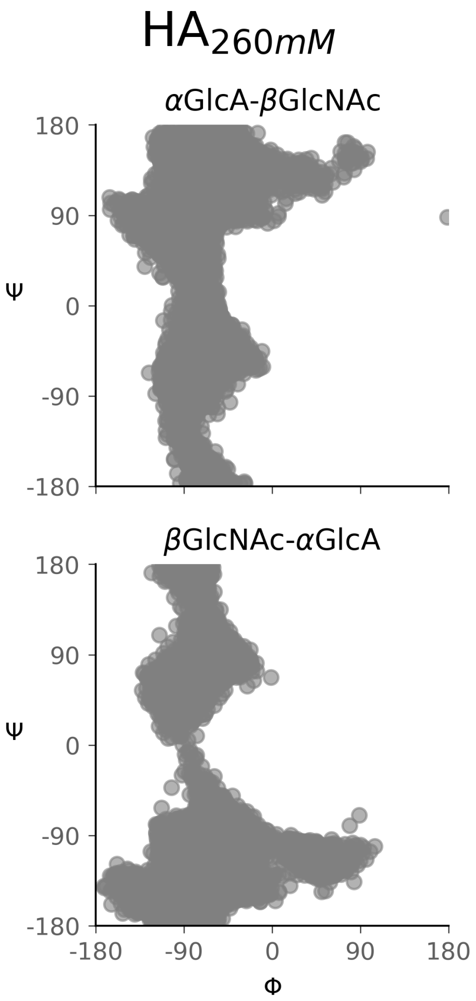


Figure S8: Distribution of the glycosidic dihedral angles $(\Phi, \Psi)$ of all the chains in $HA_{260mM}$ split by bond type.

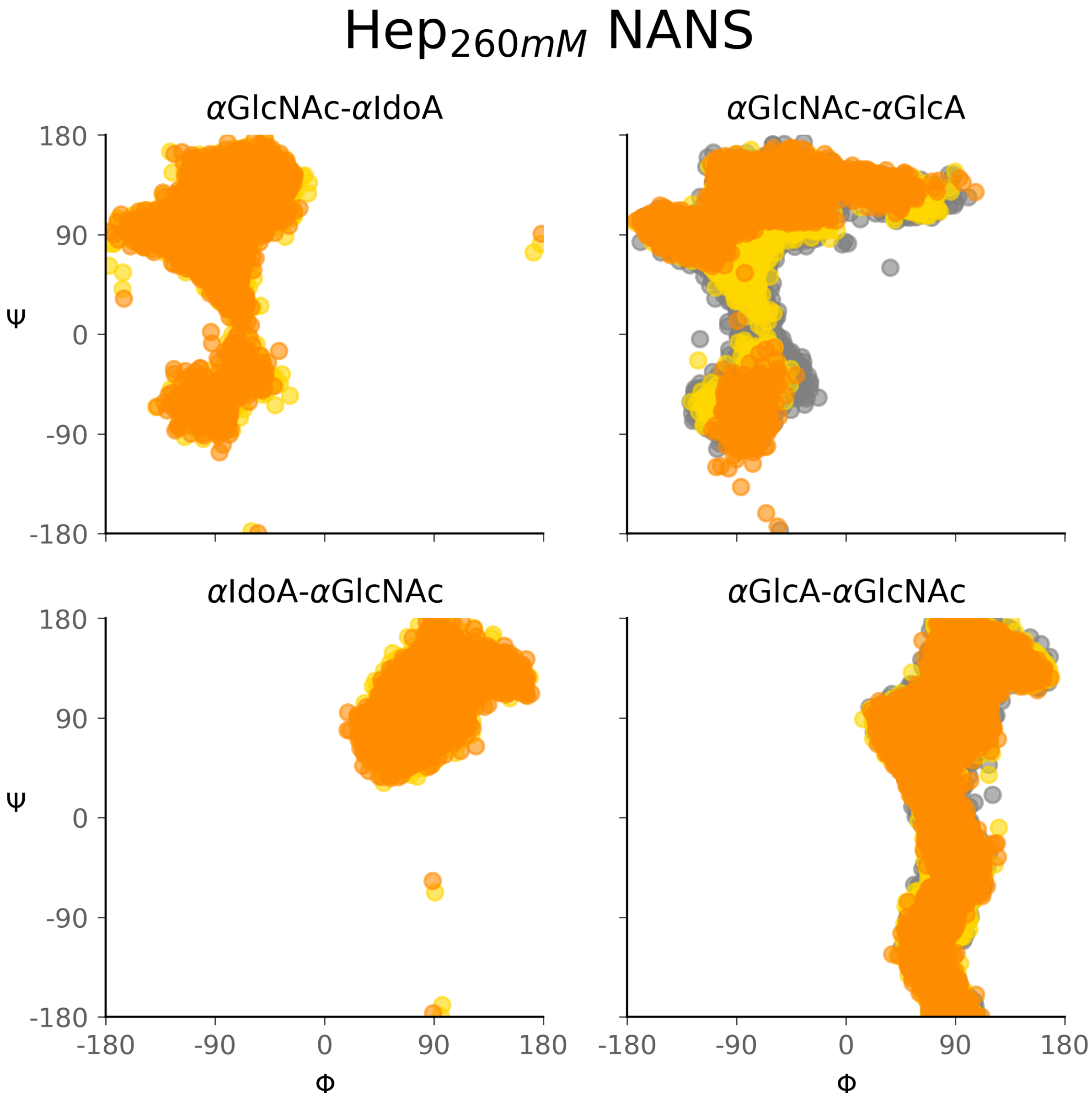


Figure S9: Distribution of the glycosidic dihedral angles $(\Phi, \Psi)$ of all the chains in $\mathrm{Hep}_{260\mathrm{mM}}$ split by bond type. Only the results for NANS chains are shown. Point color indicates the total number of sulfate groups present in the two monosaccharides forming the bond (gray: 0, yellow: 1, orange: 2).

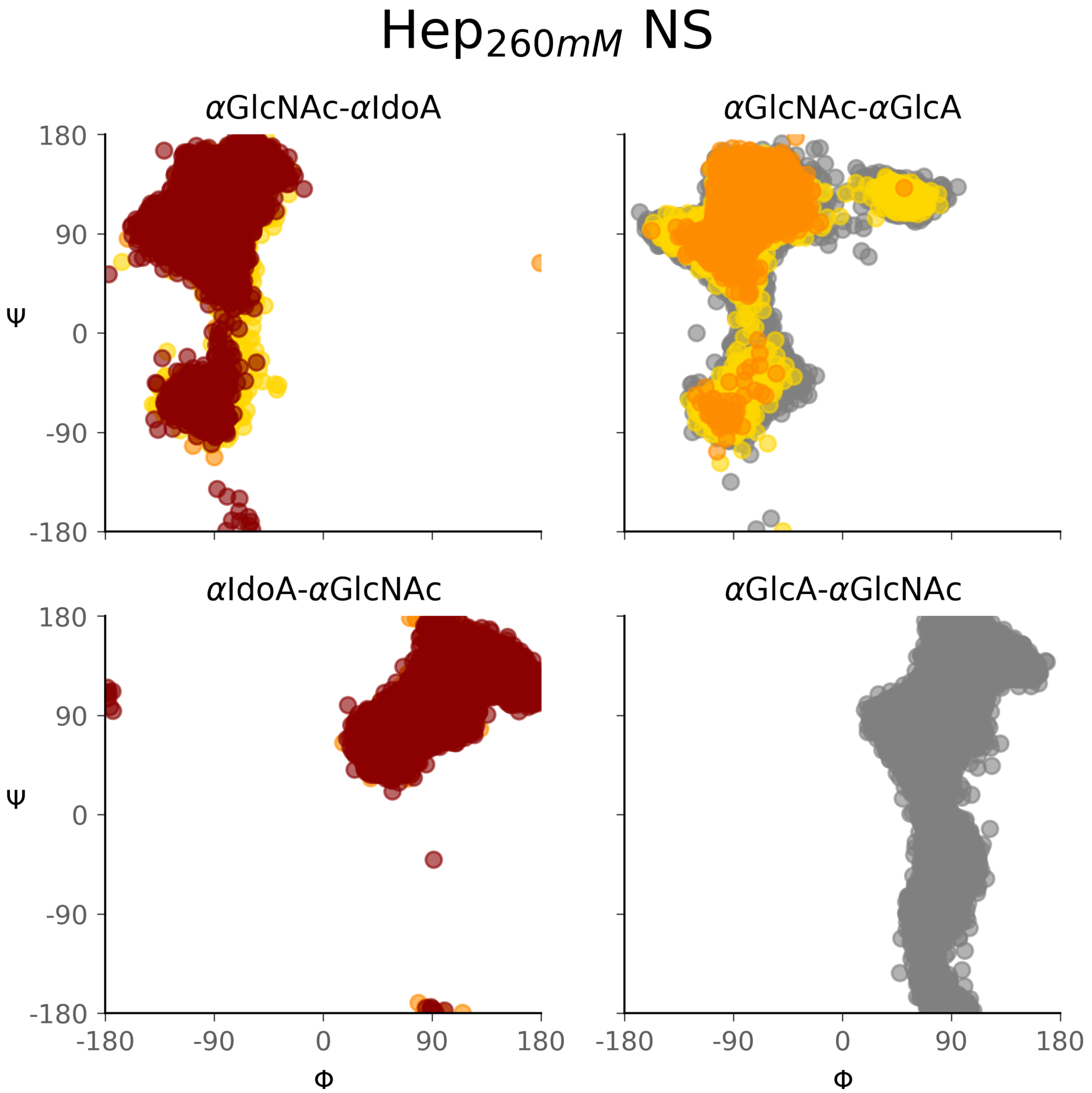


Figure S10: Distribution of the glycosidic dihedral angles $(\Phi, \Psi)$ of all the chains in $\mathrm{Hep}_{260\mathrm{mM}}$ split by bond type. Only the results for NS chains are shown. Point color indicates the total number of sulfate groups present in the two monosaccharides forming the bond (gray: 0, yellow: 1, orange: 2, red: 3).

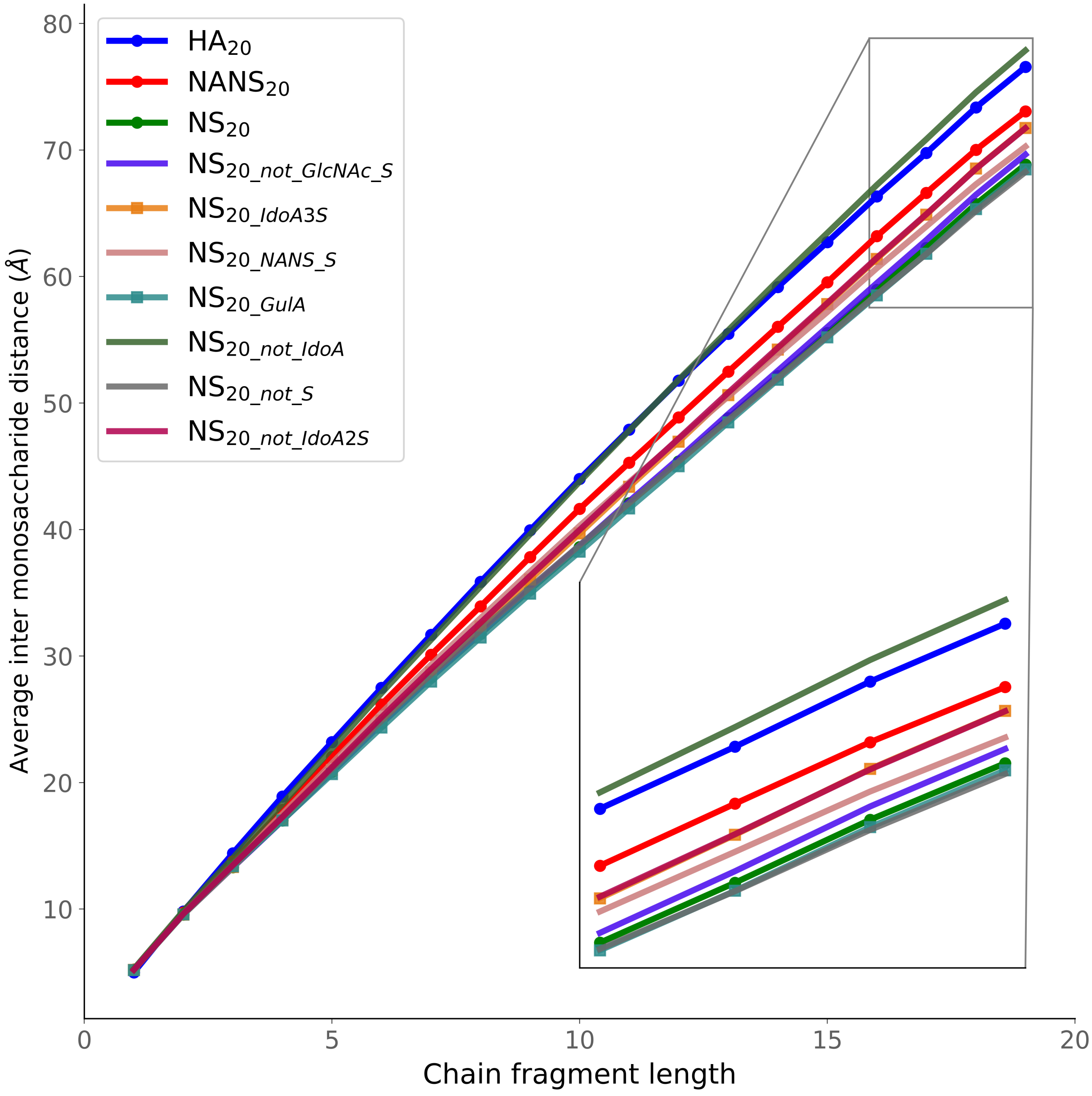


Figure S11: Average inter-monosaccharide distance as a function of the number of monomers in the 20-mer chains. The HA$_{20}$, NANS$_{20}$, and NS$_{20}$ systems are shown with dot markers since they contain the same patterns as the 50-mer simulations. NS$_{20_IdoA3S}$, and NS$_{20_GulA}$ are shown with square markers to improve visibility, as their profiles overlap with other systems.

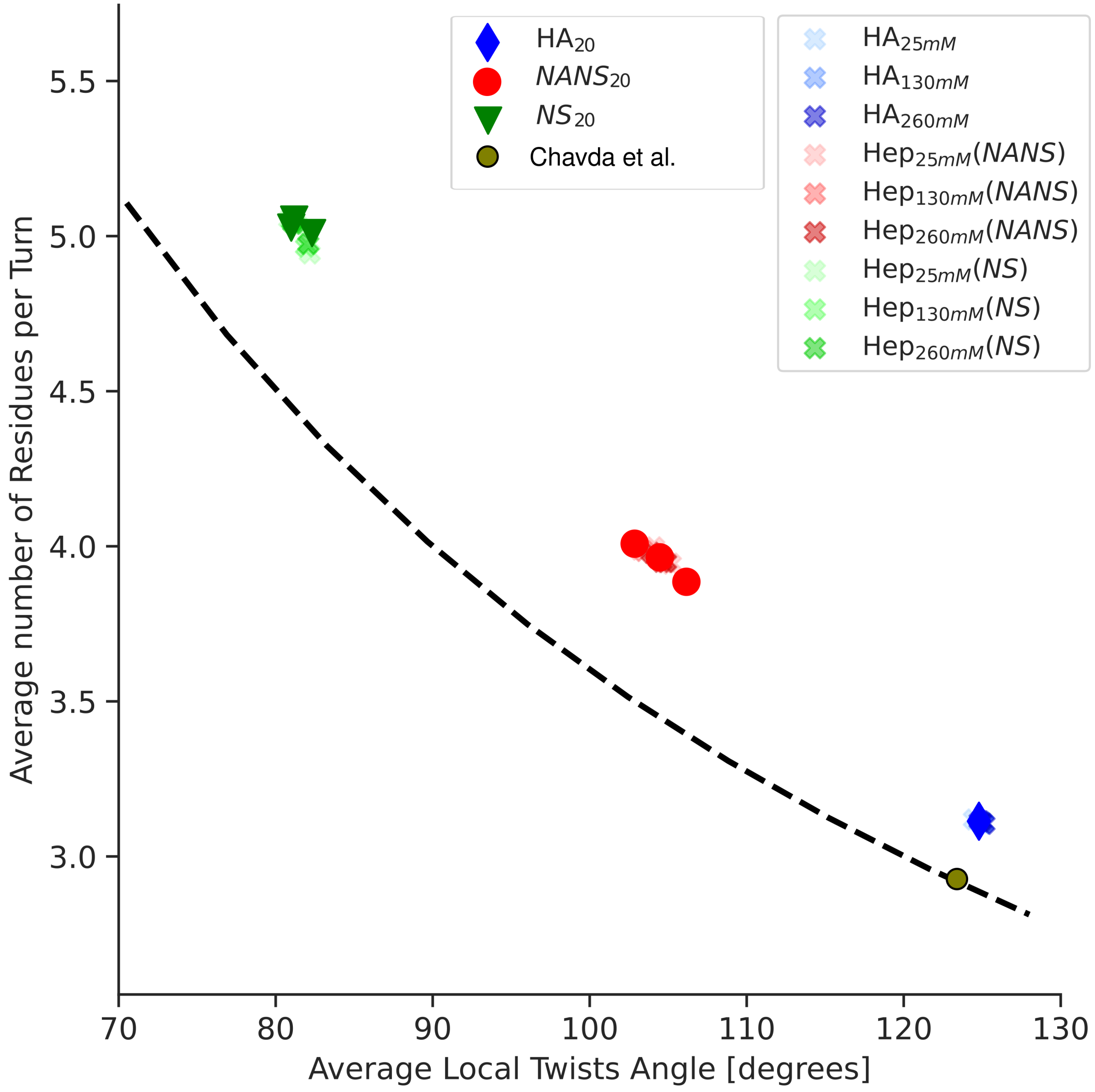


Figure S12: Helical parameters derived from the Sugeta and Tatsuo Miyazawa formalism.[S1] The data represent HA, NANS, and NS in both the 20- and 50-mer systems, as well as a structure from Chavda et al.[S2] Each point represents a sulfated domain (NS, NANS) or equivalent region in HA. The 50-mer systems have 3 points each since there are 3 sulfated regions per chain. The dashed black line corresponds to the geometric ideal helix (twist × residues per turn = 360°). The reproducibility of the metric is indicated by the fact that points from similar systems all fall within the same range.

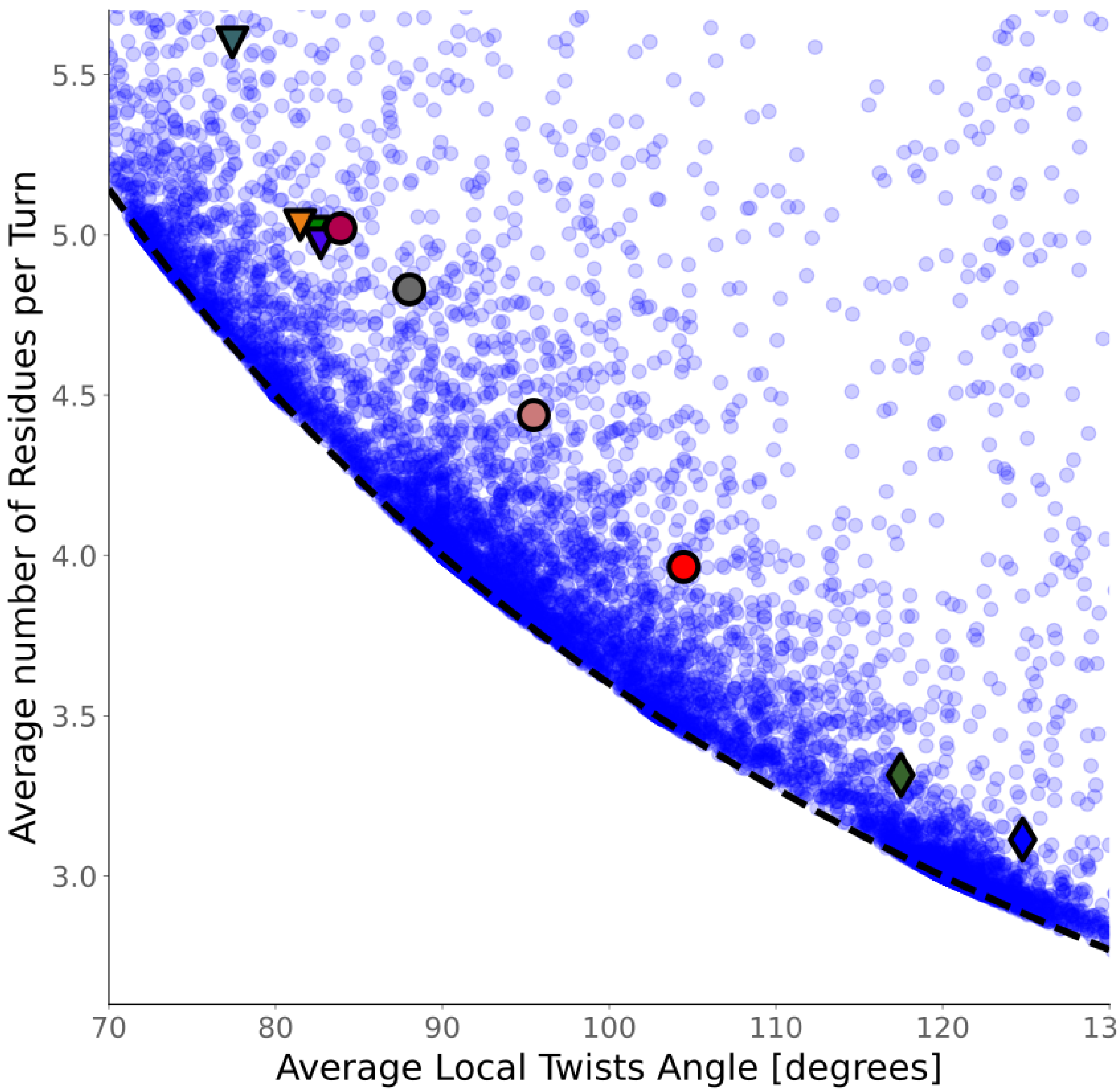


Figure S13: Average local twists and the average number of residues per turn for our custom 20-mer systems and a set of computer-generated helices with and without added noise. Semi-transparent blue points correspond to synthetic helices generated with added noise. The colored points represent the 20-mer systems following the same shape and color scheme as in Figure 6. The dashed black line corresponds to the geometric ideal helix (twist $\times$ residues per turn $= 360°$). This figure shows that all our GAG helices lie within the noisy-helix region. Therefore, the term hep–Helix should not be considered equivalent to a perfect helix but a type of secondary structure by itself.
